# Infant low-frequency EEG cortical power, cortical tracking and phase-amplitude coupling predicts language a year later

**DOI:** 10.1101/2022.11.02.514963

**Authors:** Adam Attaheri, Áine Ní Choisdealbha, Sinead Rocha, Perrine Brusini, Giovanni M. Di Liberto, Natasha Mead, Helen Olawole-Scott, Panagiotis Boutris, Samuel Gibbon, Isabel Williams, Christina Grey, Maria Alfaro e Oliveira, Carmel Brough, Sheila Flanagan, Usha Goswami

**Affiliations:** Department of Psychology, Centre for Neuroscience in Education, University of Cambridge, Cambridge, United Kingdom; Psychology and Sports Science, Anglia Ruskin University, Cambridge, United Kingdom; Department of Psychology, Goldsmiths, University of London, London, UK; Institute of Population Health, University of Liverpool, Liverpool, United Kingdom; School of Computer Science and Statistics, Trinity College Dublin, Dublin, Ireland; Laboratoire des Systèmes Perceptifs, UMR 8248, CNRS, Ecole Normale Supérieure, PSL Research University, Paris, France

## Abstract

Cortical signals have been shown to track acoustic and linguistic properties of continuous speech. This phenomenon has been measured in both children and adults, reflecting speech understanding by adults as well as cognitive functions such as attention and prediction. Furthermore, atypical low-frequency cortical tracking of speech is found in children with phonological difficulties (developmental dyslexia). Accordingly, low-frequency cortical signals may play a critical role in language acquisition. A recent investigation with infants Attaheri et al., 2022 (1) probed cortical tracking mechanisms at the ages of 4, 7 and 11 months as participants listened to sung speech. Results from temporal response function (TRF), phase-amplitude coupling (PAC) and dynamic theta-delta power (PSD) analyses indicated speech envelope tracking and stimulus-related power (PSD) for delta and theta neural signals. Furthermore, delta- and theta-driven PAC was found at all ages, with theta phases displaying stronger PAC with high-frequency amplitudes than delta. The present study tests whether these previous findings replicate in the second half of the full cohort of infants (N = 122) who were participating in this longitudinal study (first half: N=61, (1); second half: N=61). In addition to demonstrating good replication, we investigate whether cortical tracking in the first year of life predicts later language acquisition for the full cohort (122 infants recruited, 113 retained) using both infant-led and parent-estimated measures and multivariate and univariate analyses. Increased delta cortical tracking in the univariate analyses, increased ∼2Hz PSD power and stronger theta-gamma PAC in both multivariate and univariate analyses were related to better language outcomes using both infant-led and parent-estimated measures. By contrast, increased ∼4Hz PSD power in the multi-variate analyses, increased delta-beta PAC and a higher theta/delta power ratio in the multi-variate analyses were related to worse language outcomes. The data are interpreted within a “Temporal Sampling” framework for developmental language trajectories.

## 1 Introduction

Infant speech perception develops rapidly during the first year of life, yet our understanding of the neural factors underpinning language acquisition is still incomplete. Whilst infant behavioral experiments have highlighted perceptual factors contributing to early phonological and morphological development, for example perceptual ‘magnet’ effects whereby native language phonetic category prototypes pull neighbouring speech sounds towards them (2), the neural mechanisms supporting infant language acquisition are poorly understood.

Information about neural mechanisms could eventually identify early neural markers of developmental disorders of language, as neural processing is automatic and key neural mechanisms may differ in their efficiency between infants. To explore this possibility, here we contribute novel longitudinal neural data from a relatively large sample of 113 infants, utilising EEG recordings taken during the first year of life while infants listened to sung or rhythmically spoken (chanted) nursery rhymes. Nursery rhymes were chosen because of their rhythmic properties, as infant behavioural research has suggested that a sensitivity to acoustic rhythm is a universal precursor of language acquisition (3). *A priori*, it was expected that individual differences in infants’ cortical tracking of sung and chanted speech may predict later language outcomes. Accordingly, a range of neural measures are used to explore relations with subsequent language acquisition using a selection of standardised and experimental phonological, gestural and lexical measures administered between 12 to 24 months of age (4).

The auditory neuroscience of adult speech processing has revealed that accurate encoding via neuronal oscillations of amplitude modulations (AMs) nested in the envelope of the speech signal is fundamental to many different aspects of language processing (5–12). Regarding earlier development, and building on these adult insights, Temporal Sampling (TS) theory describes how the low-frequency neural oscillations that encode speech rhythm patterns may be key to learning and extracting phonological information from the speech signal (13,14). According to TS theory, auditory cortical networks operating at delta, theta, beta and gamma frequencies ‘sample’ the envelope of the speech signal at matched frequencies, thereby underpinning the phonological (linguistic) encoding of speech by infants and children. Cortical signals have been shown to track various key properties of speech, using methodological advances such as Temporal Response Functions (TRFs), an analysis technique that explores how neural signals encode continuous sensory stimuli. Adult TRF studies have revealed how cortical tracking of the speech envelope reflects both bottom-up and top-down cortical processes in adult listeners, linking to selective attention, prediction, signal parsing and speech intelligibility (7–9,15–18). Recent infant EEG data have revealed robust cortical tracking of the speech envelope (19–21), particularly in the delta and theta frequency bands (1). Indeed, patterns of cortical tracking in delta and theta frequency bands make distinct contributions to speech encoding by adults (7), and there is evidence for preferential tracking in the delta band during the first year of life (∼0.5Hz–4 Hz, see 1,22). Consequently, TS theory proposes that language acquisition by infants may depend in part on successful low-frequency cortical tracking of speech envelope information. This hypothesis regarding low-frequency cortical tracking is tested here by using TRFs.

Further, low-frequency cortical oscillations do not work in isolation from each other but show dynamic interactions during speech processing. There is a nested hierarchy of cortical oscillations that matches the frequency of multiple components of the speech signal (22). Low-frequency (delta, theta) neural phase dynamics temporally organize the amplitude of high-frequency signals in a process known as phase-amplitude coupling (PAC; (23–25)). Adult electrophysiological and MEG recordings have shown that PAC between delta-gamma, delta-beta and theta-gamma is induced in response to continuous speech (22,26,27), as well as to rhythmic speech stimuli (sung and chanted nursery rhymes, (28)). PAC is thought to group speech information into linguistic units (6,29), and accordingly could constitute another source of individual differences in language acquisition. Like natural speech, rhythmic speech induces PAC in infants and adults (1,28). Interestingly, PAC emerges as early as 4 months of age and remains stable, while both delta- and theta-band cortical tracking show developmental changes over the first year of life (1). Whilst the exact mechanisms linking PAC and language processing are still being explored in the adult literature, it is clear from this literature that PAC plays a key role in the encoding processes that are necessary for phonological learning and speech comprehension (18,22,26). Therefore, it is likely that successful PAC in infants may relate to later language outcomes.

Studies relating infant cortical tracking to later language outcomes are rare (30). However, infants at risk for dyslexia differ from not-at-risk infants in measures of acoustic processing that relate to cortical tracking, with implications for later language development (31). Meanwhile, research on the role of cortical dynamics in language development has progressed by studying older children with language difficulties. For example, recent EEG modelling studies (32) based on TS theory have shown that delta-theta PAC during passive story listening is atypical in children with developmental language disorder (DLD) while the theta/delta power ratio during passive story listening is atypical in children with developmental dyslexia. Children with dyslexia have poor phonological processing skills while children with DLD have poor syntactic skills. Based on (32), we can expect that individual differences in the theta/delta power ratio for infants may also predict subsequent language development. Furthermore, previous electrophysiological and magnetoencephalography studies testing TS theory with children with dyslexia have revealed a significant role for delta band encoding in both phonological and vocabulary development (33–36). Filtered speech interventions that enhance delta-band information also improve the theta-delta oscillatory power ratio during speech listening for dyslexic children (34). These neural data from children suggest that cortical tracking within specific low-frequency bands (i.e. delta, 0.5–4 Hz and theta, 4–8 Hz) and their interactive dynamics may be central to language acquisition by infants (13,37,38).

In the current study, we explore the potential role of individual differences in low-frequency cortical tracking, PAC and theta-delta power relations in predicting individual differences in later language acquisition. Participating infants were drawn from the Cambridge UK BabyRhythm project, a longitudinal study of 122 infants (122 recruited, 113 retained) from two- to 42-months-of-age, investigating early neural and motor tracking of auditory, visual and audiovisual rhythms in relation to language outcomes. Here we examine whether increases in low-frequency EEG power spectral density (PSD) in the delta and theta range, low-frequency cortical tracking (delta, theta and alpha; alpha band data are used as a control band, as TS theory does not accord a role to alpha-band cortical tracking regarding initial language acquisition) and PAC in response to rhythmic audiovisual speech predict performance on multiple language and communication measures administered in later infancy and toddlerhood. EEG data were collected longitudinally at 4-, 7- and 11-months in response to both nursery rhymes and during a silent state control period. The neural outcome measures were then used to predict subsequent performance in phonological, gestural and lexical measures taken from 12 to 24 months of age. It was expected that stronger cortical tracking would be predictive of better language outcomes (30). As PAC did not show development with age in (1), no strong predictions were made. However, given the DLD modelling data noted earlier (39), poorer theta-driven PAC may be linked to worse language outcomes. Further, given the dyslexia modelling data, a higher theta/delta power ratio may be linked to worse language outcomes (39). In Araújo et al. (2024), higher ratios in the dyslexic group were not driven by group differences in either delta or theta power per se, but by their joint dynamics.

## 2 Materials and Methods

### 2.1 Infant participants and ethics

All the data reported here are from the longitudinal Cambridge UK BabyRhythm project (122 infants recruited, 113 retained). The cohort was split into two halves based on recruitment date, to allow for confirmatory analyses of neural factors by comparing EEG data from the first and second halves of the cohort. The present results primarily address findings from the second half of the sample. Although results from the first half of the sample are reported in (1); it is important to highlight that minor discrepancies exist between the values presented here and those in (1). These variations stem from nuanced adjustments made to the analytical methods, elaborated upon within each respective section. Due to participant withdrawals, missed appointments and data exclusion the final number of data points for the second half of the sample was, ∼4-months (N=56), ∼7-months (N=51), ∼11-months (N=52). Missing data were due to missed appointments or technical issues with the data files, ∼4-months (N=4), ∼7-months (N=10) and ∼11-months (N=7). Other infants were excluded due to having fewer than 42 data epochs available after pre-processing, ∼4-months (N=1), ∼7-months (N=0) and ∼11-months (N=2).

The two halves of the cohort were then combined for the novel language analyses reported here (language outcome data were still being collected when Attaheri et al., 2022 (1) was being prepared). The full cohort provided EEG data for analysis as follows, ∼4-months (N=110, aged 115.5□± 5.0 days), ∼7-months (N=104, aged 213.5□±□6.1 days) and ∼11-months (N=107, aged 333.6, ± 5.0 days) [mean ± standard deviation (*SD*)].

Infants were recruited from a medium sized city in the United Kingdom and surrounding areas (recruitment period 01/10/16 to 19/12/18) via multiple means including flyers in hospitals, schools and antenatal classes, research presentations at maternity classes and online advertising. All infants were born full term (37-42 gestational weeks) and had no diagnosed developmental disorder. The study was reviewed by the Psychology Research Ethics Committee of the University of Cambridge. Parents gave written informed consent after a detailed explanation of the study and families were reminded that they could withdraw from the study at any point during the repeated appointments (8 EEG recordings at 2-, 4-, 5-, 6-, 7-, 8-, 9- and 11-months; 6 language follow-ups at 12-, 15-, 18-, 24-, 30- and 42- months). For the current report, the EEG data collected at 4, 7 and 11 months are utilized, as the nursery rhyme stimuli were only presented to the infant during these recording visits. The language outcome data are analysed using EEG measures computed from the full Cambridge UK BabyRhythm cohort (N=113) rather than separating the first and second halves of the cohort.

### 2.2 Stimuli

A selection of 18 common English language nursery rhymes were chosen as the stimuli. Audio-visual stimuli of a singing female (head only) were recorded using a Canon XA20 video camera at 1080p, 50fps and with audio at 4800 Hz. A native female speaker of British English used infant directed speech to melodically sing (for example “Mary Mary Quite Contrary”) or rhythmically chant (for nursery rhymes like “There was an old woman who lived in a shoe”) the nursery rhymes whilst listening to a 120 bpm metronome played to one ear. Although the nursery rhymes had a range of beat rates and indeed some utilized a 1 Hz rate, the metronome was used to keep the singer on time. The beat was not present on the stimulus videos, but it ensured that a consistent quasi-rhythmic production was maintained throughout the 18 nursery rhymes. To ensure natural vocalisations the nursery rhyme videos were recorded while being sung or rhythmically chanted to an alert infant.

### 2.3 EEG data collection

Infants were seated in a highchair (with neck pillow support if required) approximately one meter in front of their primary care giver, within a sound-proof acoustic chamber (on occasion it was necessary for the infant to sit on the lap of the primary care giver). EEG data were recorded at a sampling rate of 1000 Hz using a GES 300 amplifier connected to an appropriately sized 64 channel infant net (Geodesic Sensor Net, Electrical Geodesics Inc., Eugene, OR, USA). The infant was seated ∼650mm away from the presentation screen and sounds were presented at 60dB (checked by sound level meter) from speakers (Q acoustics 2020i driven by a Cambridge Audio Topaz AM5 stereo amplifier) placed either side of the screen.

Whilst the infant attended to the screen, 18 nursery rhyme videos were randomized and then played sequentially in a block, each block was then repeated 3 times (54 videos, with a presentation time of 20’ 33’’ in total). If the infant lost attention to the screen an attention grabber video was played at the end of that nursery rhyme. If the infant became too fussy a short break was allowed before continuing, otherwise the session was ended. All infants included for analysis listened to at least 2 repetitions of each nursery rhyme (minimum of 36 nursery rhymes, lasting 13’ 42’’). This stimulus period was followed by 5 minutes of silent recording (hereafter silent state). The silent state was recorded after the nursery rhymes, and commenced after a 2- minute break to check on the welfare of the participant and parent. The nursery rhymes were given first in order to maximise infant attention. Note that this may risk stimulus after-effects during the silent state (40), however the time-course of cortical tracking of the EEG signal is not comparable to fNIRs (5,41). Whilst recent studies have found oscillatory after-effects in infant EEG (42,43) such after-effects would work against our hypothesis by making it more difficult to demonstrate cortical tracking when comparing the nursery rhyme data to the silent state data, making the PSD and TRF analyses more stringent. To ensure compliance from the infant, and therefore a good EEG signal during the silent state, it was sometimes necessary for a researcher to sit alongside the infant. To ensure consistency across participants, the researcher performed the same action of silently blowing bubbles and showing the same picture book to the infant during this period.

### 2.4 EEG preprocessing

All analyses were conducted with custom-made scripts in Matlab 2017a (The MathWorks, Inc., Natick, MA) incorporating the EEGLab toolbox (44). The EEG data recorded in response to the nursery rhymes and the silent state were treated as one continual data set for the initial preprocessing steps. Therefore, they underwent the same cleaning and filtering procedure prior to epoching. The 60 EEG channels (present on the infant-sized EGI Geodesic sensor nets) were filtered using the pop_eegfiltnew function from the EEGLab toolbox. The data were either filtered into a broadband signal for PSD and PAC analysis (0.5–45 Hz, methods for later filtering into PAC bands detailed in section 2.6) or into specific frequency bands for the mTRF analyses: delta (0.5–4 Hz), theta (4–8 Hz), or alpha (control band, 8–12 Hz). A zero-phase bandpass Hamming-windowed FIR filter was used, with transition band widths of 2 Hz and cut-off frequencies at -6 dB (0-46 Hz, 0–5 Hz, 3–9 Hz, and 7–13 Hz, respectively). Next, the EEG data were then down sampled to 100 Hz to reduce computational load. Artifact Subspace Reconstruction (ASR; *clean_asr* EEGLab function (44,45)) was used separately on each of the data sets to clean noise artifacts from the data by identifying and removing bad principal components via a modified PCA procedure (see (1)). ASR effectively removed the transient or large-amplitude artifacts common in infant EEG data. Further bad channels were identified via probability and kurtosis and were interpolated (via spherical interpolation). A channel was rejected and interpolated if it was 3*SD* away from the mean (average number of interpolated channels; 4mo = 6.9, 7mo = 6.5, 11mo = 6.6). All channels were re-referenced to a 60-channel average reference. Next, the EEG responses to the 18 nursery rhymes were epoched into trials aligned to the start of a nursery rhyme phrase (e.g. “Mary had a little lamb”). Each nursery rhyme contained approximately 5 phrases (*M number of phrases per nursery rhyme* ± *SD*: *4.61 phrases ±4.22,* range =1:20) producing EEG responses to 83 phrases (*M length* ± *SD*: 4.23sec ±0.88) which were repeated a maximum of 3 times in the experiment (249 epochs in total). This epoching procedure was selected as infant EEG typically contains short, irregular, movement artifacts and using shorter epochs increases the likelihood of generating a whole artifact-free epoch whilst maintaining long enough epochs for optimising the model fit with the mTRF toolbox (46). The infants occasionally rested their head or touched the EEG cap in such a way that specific channels showed short bursts of noise whilst the rest of the channels remained clean. To account for this, and retain a larger volume of data, a second stage of channel rejection and interpolation was conducted epoch by epoch on each of the four filtered datasets (broadband: 0.5–45 Hz, delta: 0.5–4 Hz, theta: 4–8 Hz, and alpha: 8–12 Hz). Whilst this second round of epoch by epoch channel rejection and interpolation varies slightly from classical EEG pre-processing, it was considered a legitimate trade off to retain data from clean channels whilst removing short, irregular, movement artifacts present only in a few channels. Per epoch, probability and kurtosis were used to identify bad channels and were interpolated (via spherical interpolation) if they were 3*SD* away from the mean. Finally, bad epochs were rejected with the *pop_autorej* function (EEGLab), removing epochs with fluctuations above 1000uV and values outside a 3*SD* of the probability threshold. Due to the interpolation and ASR procedure, less than 1% of the epochs were rejected via *pop_autorej*. This step was maintained as a sanity check to ensure all extreme fluctuations had been removed. Due to the increased susceptibility of PSD and PAC analyses to noise, manual rejection was conducted, on the 0.5-45 Hz data set, to remove noisy periods of data from both the silent state and nursery rhyme data prior to running the PSD and PAC analyses. To minimize within trial data discontinuities manual rejection boundaries were limited to in-between phrases. The separation of the nursery rhyme and silent state EEG data occurred after this manual rejection so that the PSD analysis could be conducted separately for the two conditions.

### 2.5 mTRF auditory stimuli preprocessing

The envelope of the auditory signal was extracted by taking the absolute value of the analytic signal generated by the Hilbert transform (Matlab). As the lower frequencies of the envelope are linearly relatable to the EEG signal (10,11,47) the envelope of the stimuli was filtered between 0.5 Hz and 15 Hz (lowpass; 6^th^ order Butterworth filter. Highpass; 9^th^ order Butterworth filter) and this was used for all individual band comparisons. The resultant envelopes were normalised using *nt_normcol* (Noisetools). Finally, the stimulus envelopes were down sampled to 100 Hz to match the EEG signal. The preprocessed and filtered envelopes, from now on will be referred to as envelopes.

### 2.6 Multivariate Temporal Response Function (mTRF)

TRFs describe the linear relationship between an input and an output signal, taking into account that such a relationship may not be instantaneous and could extend over a certain window of time (46). TRFs can be estimated with system identification methodologies, such as the multivariate temporal response function (mTRF), which is a de-convolution method based on multiple lagged linear regression. Here, we applied the mTRF (46) in the backward direction to reconstruct the stimulus envelope from the EEG signals. In turn, this informs us on the strength of the stimulus-EEG relationship, which we take as an index of cortical tracking of the stimulus envelope. The backwards ‘stimulus reconstruction’ mTRF model has important advantages compared to the forward linear encoding model which has previously been used in infant studies (19,20). Specifically, backward TRFs combine data from all the EEG channels simultaneously in a multivariate manner to reconstruct the univariate stimulus envelope, while forward TRF models only predict one EEG channel at a time.

After preprocessing, the EEG responses to each of the nursery rhymes were averaged (data across a maximum of three repeats of the 83 nursery rhyme phrases were averaged), thereby creating 83 “averaged trials”. This averaging was conducted to improve the signal to noise ratio of the data for the mTRF. The strength of the cortical tracking in each EEG frequency band was evaluated using leave-one-out cross-validation per infant (46), separately for each frequency band, delta (0.5-4Hz), theta (4-8Hz) or alpha (8-12Hz). First, the average EEG trial epochs (up to 83) were normalized using the function nt_normcol (Noisetools, http://audition.ens.fr/adc/NoiseTools/). The normalized trials were then rotated M-1 times, with each rotation serving once as the test set while the rest formed the training set. The averaged training models were used to reconstruct the stimulus by convolving with the test data. Pearson’s correlation (r) measured the correlation between the reconstructed and original stimulus envelopes. This process was repeated for all M-1 rotations, and an individual’s average r value was computed to prevent overfitting. The average r value represented how well the model reconstructed the stimulus envelope, when trained with either the delta (0.5-4Hz), theta (4-8Hz) or alpha (8-12Hz) EEG band data. Chance r scores were created for each participant to measure the average stimulus reconstruction (r) that could be obtained by chance. The chance TRF procedure was conducted per participant for each frequency band (one shuffled model per participant). To obtain a chance permutation of the data, whilst maintaining phase integrity, each of the stimulus envelopes were first reversed and a random circular shift was applied. Next, the mTRF cross-validation was run in the same way as the real data (see above for details), to give a stimulus reconstruction (r) value. The *r* value using the shuffled model was used as the chance *r* score per participant. Please note that the chance *r* values in Table 4 and Figure 2, for the first half of the sample, differ slightly from (1) as a single random model per participant rather than an average random model was implemented here. This test aims to determine if the average value across participants is above “chance”. This is a case of subsampling in which the resulting distribution should match the distribution of the population. A linear mixed effects model (LMEM) was next constructed using the same parameters as in Attaheri et al., 2022 (1), to discover if the mean of the “real” model was larger than the mean of the “shuffled” model, across participants.

### 2.7 Phase amplitude coupling (PAC)

To assess PAC, a modified version of the WinPACT plugin (EEGLab) (44) was used to acquire normalised modulation index (nMI) values (48) a measure adapted from Canolty and colleagues’ modulation index (MI) (23). This measure is a widely validated metric of the coupling strength and preferred phase between two frequencies. For each infant’s EEG data, low-frequency phase (LFP) and high-frequency amplitude (HFA) were extracted with a zero-phase FIR filter (Hamming window), separately for all 60 electrodes, from the pre-processed 0.5-45 Hz data. LFP data were filtered in 1 Hz steps from 2 to 8 Hz. Each centre frequency had a 2 Hz filter bandwidth around the centre frequency. HFA centre frequencies were extracted from 17.5 Hz to 42.5 Hz, in 5 Hz steps with a 5 Hz filter bandwidth around the centre frequency, Next, a normalised modulation index value was calculated for each channel using a 5 second sliding analysis window (with 2.5 second overlaps)(23,48). To identify windows displaying significant PAC, 200 surrogate statistical iterations were created for each window. A statistically normalised MI estimate was obtained for each analysis window by subtracting the mean and dividing by the standard deviation obtained from a Gaussian fit of surrogate MI estimates (nMI = (Canolty’s MI – surrogate MI Mean) / surrogate MI Std). This statistical procedure is based on the work of Özkurt and Schnitzler, 2011 (48) and implemented via the winPACT EEGLab plugin. Each iteration of the surrogate data was created by shuffling the high-frequency amplitude time series via circular rotation. A MI estimate was obtained for each of the 200 surrogate data iterations, from which a 95% confidence interval (CI) was calculated. This step accounted for the mean and standard deviation of the surrogate data set, thus creating an appropriate threshold for the frequency band analysed. Finally, the 95% CIs were averaged across all windows to generate a statistical threshold for each channel per participant. All the nMI windows that passed the 95% CI significance test were averaged per channel for each of the PAC pairs (i.e. each LFP and HFA step) separately for each participant. The channel exhibiting the strongest MI, within predefined phase and amplitude band groupings (delta/beta, delta/gamma, theta/beta, theta/gamma, with delta=2-4Hz, theta=4-8Hz, beta=15-30Hz and gamma=30-45Hz), was taken forward to use as the candidate predictor for the LMEM and for the group level grand average plots. Due to the Nyquist frequency required for our 5 second window size, the first frequency band analysed in the PAC was 2Hz. See (1) for a detailed description of the PAC method and analysis.

### 2.8 Spectral analysis (periodogram PSD estimate)

A one-sided PSD estimate was conducted separately for each channel of the pre-processed 0.5-45 Hz data using the periodogram function (Matlab). The grand average PSD across channels was next calculated by averaging across channels, recording type (nursery rhymes [NR], silent state [SS]) and ages (4-, 7- or 11-months). Visual inspection revealed three frequency peaks centered around 1.92 Hz, 4.05 Hz and 4.35 Hz (Fig. S1). The spectra for each individual age group per recording type were next calculated and taken forward for further analysis (Fig. 1). To allow for individual participant variation, the maximum value within a 0.25Hz window centered on the peak of interest was taken per participant for further analysis. A smaller window was used around the peaks of interest than used in Attaheri et al., 2022 (1), this was necessary to avoid overlap between the 4.05 Hz and 4.35 Hz windows. See (1) for a detailed description of the PSD method and analysis. We acknowledge that manual rejection may have introduced some discontinuities, and we took steps to minimize their effect on the PSD analysis. First, a relatively large amount of data was used in the periodogram to achieve strong frequency resolution. Second, zero-padding was applied to ensure that all data files, regardless of the amount of manual rejection, were the same size, resulting in an equal number of frequency bins across participants. Additionally, a Hamming window was used to taper the edges of the data, smoothly reducing the signal to zero at the boundaries and minimizing the impact of discontinuities. This tapering improves frequency resolution and reduces spectral leakage. Overall, the combination of manual rejection and these mitigation steps was considered a valid approach to removing noise that could compromise the PSD analysis.

**Figure 1.**
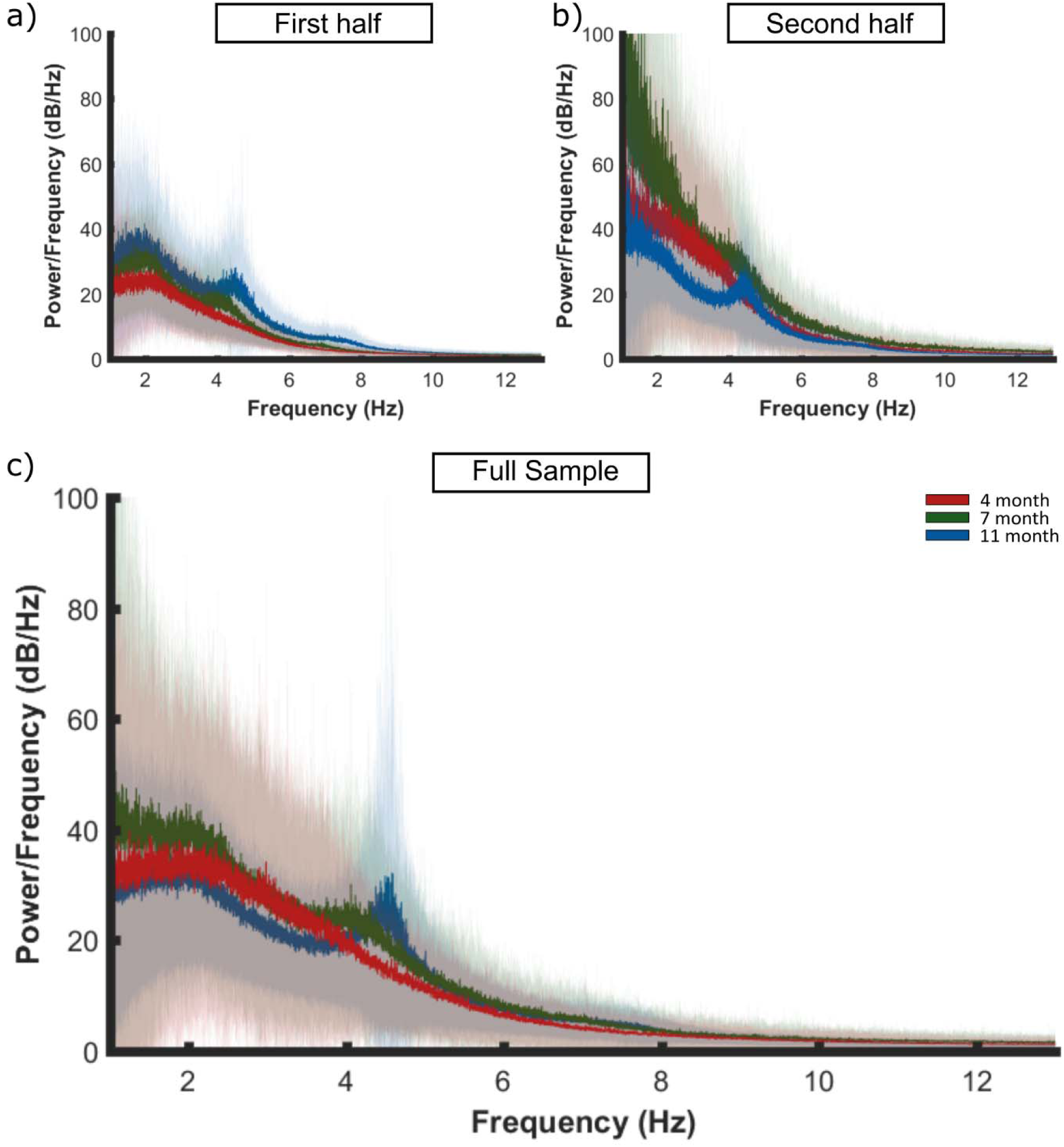
Spectral decomposition of the EEG signal averaged across the a) first half, b) second half and c) full infant cohort (1–12 Hz plotted, from 0.5 to 45 Hz calculated), in response to nursery rhyme stimulation. A periodogram was used to obtain a power spectral density (PSD) estimate separately for 4- (red), 7- (green) and 11- (blue) months data. First the PSD estimate was averaged across channels per participant. Bold lines indicate the grand average mean values and pale shading plots the standard deviation of the data across the participants. Outlier analysis was also conducted to remove extreme data points that would compromise the LMEM. Panel A is reproduced from Attaheri et al. 2022 (1), with permission.

#### 2.8.1 Theta / delta (PSD) ratio

The PSD peak per participant at 4.35 Hz and 1.92 Hz, taken from the spectral decomposition of the EEG signal described above, was used to create a theta / delta ratio calculation. Per participant, the 4.35 Hz PSD value was divided by the 1.92 Hz PSD value (4.35/1.92) to give a theta / delta ratio.

### 2.9 Language measures

A full description of the language measures used here can be found in Rocha et al., (4). The parent-estimated measures were taken from a carer-completed questionnaire, the UK-CDI. The UK-CDI estimates both receptive and productive vocabulary (49), which we refer to here as language comprehension or language production. In the current report the parent-estimated 24- month CDI measures were selected for analysis, as by the age of 2 years both word comprehension and word production data showed a wide range of scores, enabling more sensitive measurement of individual differences (see (4), for detailed rationale). Only measures up to the age of 24 months are included here, as the 30- and 42-month measures are still being processed. Five measures were selected as the infant-led (in that the infant made an active response) language measures. These were (i) gesture (adapted from (50)), in which pointing towards a present and absent target was used as a pre-verbal measure of communicative intent at 12 months, and (ii) the computerized comprehension task (CCT), a touch screen-based measure of infant vocabulary administered at 18 months, in which the infant is asked to select a picture of a named item (51). Thirdly a non-word repetition (NWR) task based on (52), was used to measure the child’s productive phonology at 24 months and generate three different measures: (iii) accurate reproduction of consonants, (iv) number of syllables reproduced, and (v) stress patterns produced (canonical or non-canonical, see Rocha et al., (4)). The NWR task measured toddlers’ phonological production of both nonsense words (e.g. “punky”) and real words (e.g. “puppy”) via a game about naming a series of toys.

### 2.10 Language outcomes: statistics

Our statistical approach was decided *a priori* and was established when exploring the language outcome measures (4). We ran a multivariate model for each of the infant-led and parent-estimated groups of measures. Given that missing cases vary across the multiple tasks, and that the multivariate models employed use list wise deletion, additional follow-up univariate models for each of the predictors were run separately, to utilise the breadth of data collected and maximise statistical power. This approach was applied across multiple BabyRhythm studies using different neural measures as language predictors (see (53), using phase measures, and (54), which uses neural responses to visual-only speech), accordingly an alpha threshold of 0.05 was applied for the multivariate models. As the follow-up univariate analyses required multiple comparisons, one per language measure, they were Bonferroni-corrected for the number of language measures in the group (i.e. 5 for infant-led, and 2 for parent-estimated). Respective alpha levels were therefore 0.01 and 0.025. Although the main multivariate tests may show that some predictors had a significant effect and others did not, we did not remove any of the original covariates in the univariate models, to ensure comparability between the multivariate and univariate tests.

Due to the inherent differences between the infant-led and parent-estimated measures, separate multivariate linear models were conducted for each, providing a global measure of language abilities taken from either infant-led or parent-estimated language measures. Multivariate linear models (using Rstudio *lm* function) were run separately for the PSD, mTRF, theta-delta ratio and PAC measures. Secondly, Bonferroni-corrected individual univariate linear models were run, which typically included more infants, in order to explore the contribution of the early neural measures to the individual language measures.

## 3 Results

### 3.1 PSD, mTRF, PAC and theta/delta ratio

First, we present the power spectral density (using a periodogram; PSD), cortical tracking (using multivariate temporal response functions; mTRFs), phase-amplitude coupling (normalised modulation index, nMI; PAC) and theta/delta PSD ratio analyses using EEG recorded from the second half of our cohort of infants. We compare these data to the analyses from the first half of the cohort (1), and also merge the neural data to explore these neural factors in the combined full sample of 113 infants. Due to missed recording sessions, technical issues with data files and not enough trials after preprocessing, less than 113 data points are included in each analysis. Furthermore, outlier analysis was also conducted to remove extreme data points that would compromise each of the LMEMs. The number of data points included are given separately for each analysis.

#### 3.1.1 Power spectral density (PSD) analyses

The distribution of low-frequency oscillations in the EEG recorded from the second half of the sample and the full cohort were obtained using the power spectral density (PSD) estimate (Fig. 1). The results from the first half of the cohort (1) are reproduced for comparison. Given the differences in the location of the PSD peaks for the first and second halves of the cohort, the distribution of PSD peaks in low-frequency power for the full sample was first established by creating a grand average PSD estimate across channels, recording type (nursery rhymes, silent state) and ages. Visual inspection revealed three frequency peaks centered around 1.92 Hz, 4.05 Hz and 4.35 Hz (Fig. S1, and in the second half of the sample, Fig. S2). The distribution of the PSD peaks were similar to those observed in the first half of the sample (see (1) and Fig. S3). It is important to note that the PSD peaks matched the most prominent modulation spectrum peaks that were present in the nursery rhyme stimuli (see Fig. S4–S6), which were respectively within the delta and theta band definitions used for the EEG analyses (delta, 0.5–4 Hz; theta, 4–8 Hz).

Whilst infant frequency bandings appear to change throughout infancy (55–58), there are no prior infant data regarding center frequencies in studies using speech stimuli. Therefore, we selected the delta and theta boundaries based on the prior adult speech processing literature, and this decision is supported by the modulation bands identified by modelling the temporal modulation architecture of the 18 nursery rhymes used in our study (59)). Further, when the EEG from the full cohort was analysed, the ∼4Hz peak split into two locations, peaking at 4.05 Hz and 4.35 Hz. Both theta peaks were therefore retained for further investigation. Whilst the relevant data are provided in the results text, the mean and standard error of the PSD peak amplitudes for both silent state and nursery rhyme recording conditions, are also provided in Table S1. Plots of the silent state PSD are also provided to aid comparison (Fig. S7).

To investigate whether the PSD was significantly higher in response to the nursery rhymes compared to the silent state, and whether this relationship was affected by the age of the participants, a linear mixed effects model (LMEM) was conducted separately for the 1.92 Hz, 4.05 Hz and 4.35 Hz peaks. A random intercept (AR1) per participant was included to account for individual differences across the 3 recording sessions (4-, 7- and 11-months). The full sample submitted to the LMEM was as follows, 4-months (N=96), 7-months (N=88) and 11-months (N=92). This process was conducted separately for the first half of the sample, the second half of the sample and the full cohort (Table 1). To allow for individual variation across participants, a maximum peak value per participant was taken from the participants’ 60 channel averaged data, within 0.25 Hz windows centered around the peak of interest (1.92 Hz, 4.05 Hz and 4.35 Hz). Note that a smaller window was used around the peaks of interest than used in Attaheri et al., 2022 (1), this was necessary to avoid overlap between the 4.05 Hz and 4.35 Hz windows. Fixed effects of recording type (nursery rhymes, silent state), age (4-, 7- or 11-months) and a recording type by age interaction were included in the model. The ∼4.35 Hz peak is reported first, as this was the most robust.

**Table 1.**
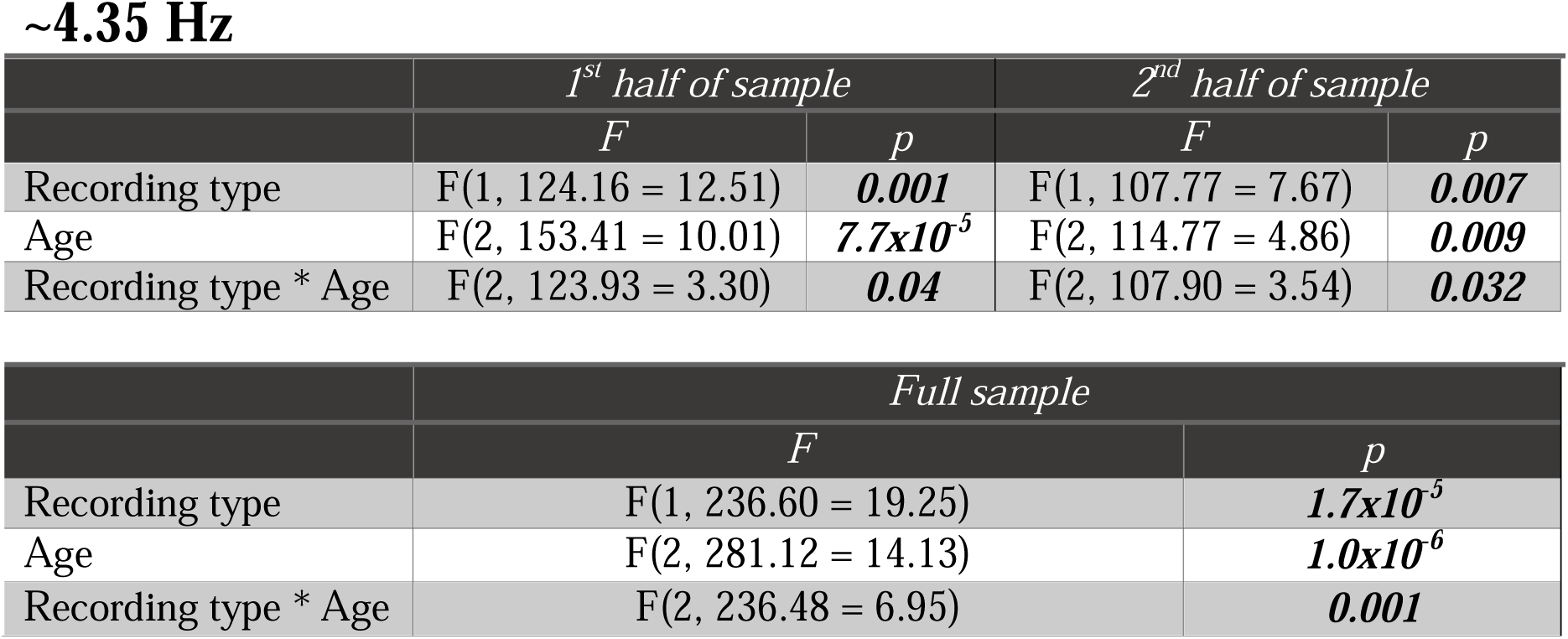
Tests of fixed effects for LMEM conducted on PSD power peak at 4.35Hz. Results reported for separate LMEMs run on the first half, second half and full sample of data. Significant results (p<0.05) denoted by ***bold italic text***. The Satterthwaite approximation was applied to approximate the degrees of freedom, due to the missing cases.

~4.35 Hz
|  | *1<sup>st</sup> half of sample* |  | *2<sup>nd</sup> half of sample* |  |
| --- | --- | --- | --- | --- |
|  | *F* | *p* | *F* | *p* |
| Recording type | F(1, 124.16 = 12.51) | **0.001** | F(1, 107.77 = 7.67) | **0.007** |
| Age | F(2, 153.41 = 10.01) | **$7.7 \times 10^{-5}$** | F(2, 114.77 = 4.86) | **0.009** |
| Recording type * Age | F(2, 123.93 = 3.30) | **0.04** | F(2, 107.90 = 3.54) | **0.032** |

Table 1. Tests of fixed effects for LMEM conducted on PSD power peak at 4.35Hz. Results reported for separate LMEMs run on the first half, second half and full sample of data.
|  | *Full sample* |  |
| --- | --- | --- |
|  | *F* | *p* |
| Recording type | F(1, 236.60 = 19.25) | **$1.7 \times 10^{-5}$** |
| Age | F(2, 281.12 = 14.13) | **$1.0 \times 10^{-6}$** |
| Recording type * Age | F(2, 236.48 = 6.95) | **0.001** |

The 4.35 Hz peak showed the largest stimulus-related increase in PSD of the three peaks analysed. The LMEM’s revealed a significant effect of recording type, with the nursery rhyme (NR) condition producing significantly greater PSD power than the silent state (SS; Table 1). A significant effect of age was observed, with ∼4.35 Hz power increasing parametrically with age from 4- months to 7- months to 11- months (estimates of fixed effects provided in Table S2). The observed recording type by age interaction was due to a developmental increase in NR induced power above the SS. Bonferroni-corrected pairwise comparisons showed that PSD increased significantly between 4- and 11-months (M increase = 18.41, SE ± 7.43, *p* = 0.043) as well as between 4- and 7- months (M increase = 30.56, SE ± 5.77, *p* = 6.7×10^-7^). Hence there was a developmental increase in 4.35 Hz PSD for both the nursery rhymes and the silent state. This may indicate age-related maturation of theta activity in general. Importantly, we also observed a stimulus-driven interaction with age for the 4.35 Hz PSD peaks, with the largest increase observed at 11-months (4-months, NR, *M* = 38.9 *SE* ± 4.9; SS, *M* = 37.9 *SE* ± 5.5. 7-months, NR, *M* = 63.0 *SE* ± 6.9; SS, *M* = 50.6 *SE* ± 7.2. 11-months, NR, *M* =84.8 *SE* ± 5.8; SS, *M* = 53.1 *SE* ± 6.0). This showed that the interaction was driven by a greater PSD in response to the nursery rhymes at 11-months. This pattern replicated the theta PSD findings observed in the first half of the sample (see (1)). In summary, the peak at 4.35 Hz showed robust replication across the first and second half of the sample, with consistent increases in NR related power displaying a developmental increase in PSD magnitudes, above SS, indicating a highly reliable peak.

For the 4.05 HZ peak, the LMEM results replicated less well across the first half of the sample, the second half of the sample and the full cohort. Significant effects of both recording type and age were only observed when analysing the full cohort. The significant effect of age was driven by the 7-month (*M* = 63.1 *SE* ± 6.0) PSD power being larger than the 11-month (*M* = 52.5 *SE* ± 3.2), however this result is in part due to the 0.25 Hz window not fully encapsulating the 11-month peak for all participants (Fig. 1). A significant effect of recording type was observed in the second half of the sample, with a trend (p<0.10) observed in the first half of the sample (Table 2). A significant effect of age was observed in the first half of the sample with a trend (p<0.10) towards significance observed in the second half of the sample. As the data were more robust for the 4.35 HZ peak, and as the 0.25 Hz window did not fully capture the 4.05 Hz peak for all participants, the 4.35 Hz PSD peak was taken forward as the ‘theta’ peak for analyses with the language data.

**Table 2.**
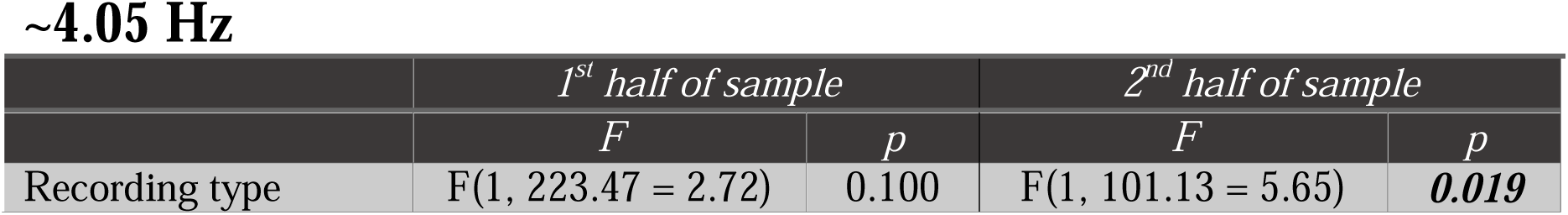
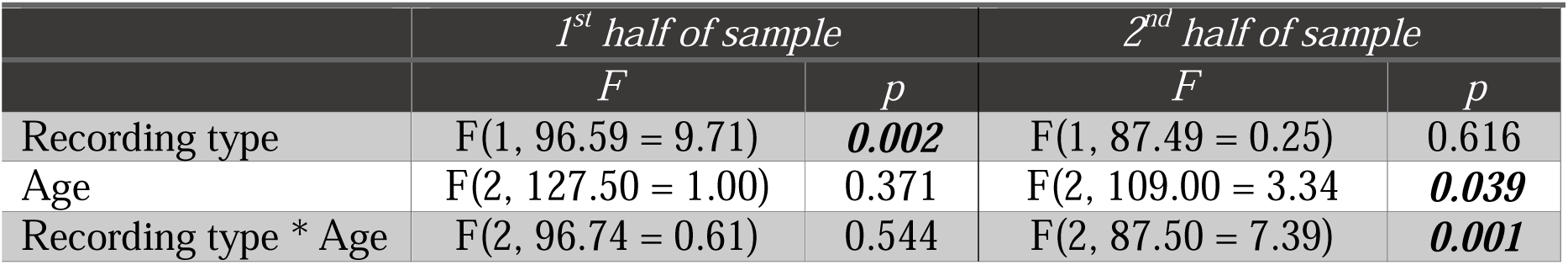
Tests of fixed effects for LMEM on PSD power peak at 4.05Hz. Results reported for separate LMEMs run on the first half, second half and full sample of data. Significant results (p<0.05) denoted by ***bold italic text***. The Satterthwaite approximation was applied to approximate the degrees of freedom, due to the missing cases.

For the 1.92 Hz peak, the LMEM conducted on the full cohort revealed a significant effect of recording type (Table 3). However, there were differences in this effect across the first and second halves of the sample. Upon further inspection, the non-significant level of recording type in the second half of the sample was due to the large amount of PSD power in the silent state (SS) data at 4-months (4-month SS, *M* = 116.3 *SE ± 10.6,* NR, M= 97.3 *SE ± 10.2*). Posthoc paired sample t-tests on the 7- and 11-month data, for the second half of the sample, both showed a consistent stimulus-induced increase in PSD power, matching the first half of the sample (7- month SS, *M*= 116.3 *SE ±13.5,* NR, *M*= 124.2 *SE ±13.1, p* = 0.007; 11-month SS, *M=76.7 SE ± 11.9 ,* NR, *M*= 94.0 *SE ± 11.6 , p* = 1.90×10^-27^). A significant recording type by age interaction was observed in both the second half of the sample and the full cohort. In conclusion, the 1.92Hz stimulus-related increase in PSD power is broadly replicated across the first and second halves of the sample once the anomalous SS power, present in the 4-month data in the second half of the sample, was taken into account.

**Table 3.**
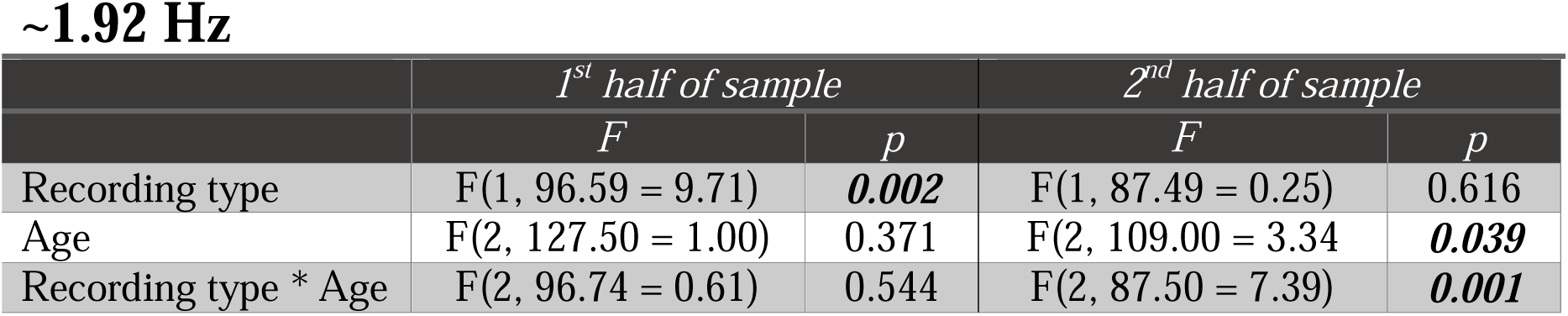
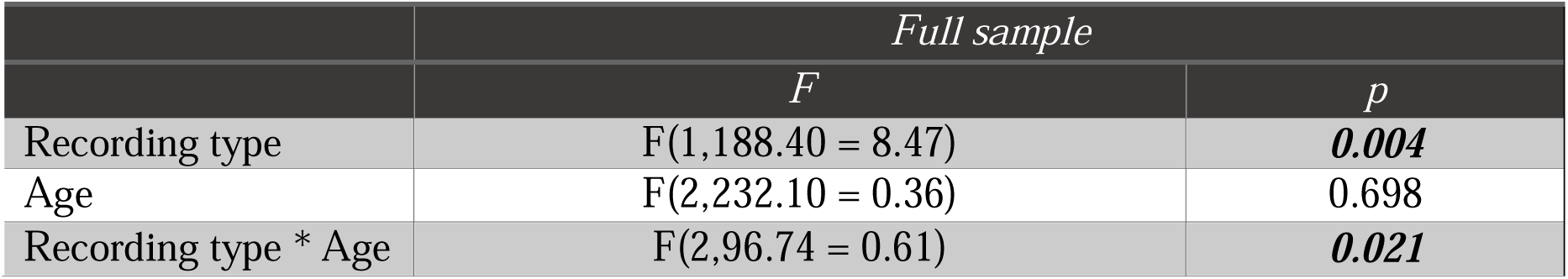
Tests of fixed effects for LMEM on PSD power peak at 1.92Hz. Results reported for separate LMEMs run on the first half, second half and full sample of data. Significant results (p<0.05) denoted by ***bold italic text***. The Satterthwaite approximation was applied to approximate the degrees of freedom, due to the missing cases.

~1.92 Hz
|  | *1<sup>st</sup> half of sample* |  | *2<sup>nd</sup> half of sample* |  |
| --- | --- | --- | --- | --- |
|  | *F* | *p* | *F* | *p* |
| Recording type | F(1, 96.59 = 9.71) | **0.002** | F(1, 87.49 = 0.25) | 0.616 |
| Age | F(2, 127.50 = 1.00) | 0.371 | F(2, 109.00 = 3.34) | **0.039** |
| Recording type * Age | F(2, 96.74 = 0.61) | 0.544 | F(2, 87.50 = 7.39) | **0.001** |

Table 3. Tests of fixed effects for LMEM on PSD power peak at 1.92Hz.
|  | *Full sample* |  |
| --- | --- | --- |
|  | *F* | *p* |
| Recording type | F(1,188.40 = 8.47) | **0.004** |
| Age | F(2,232.10 = 0.36) | 0.698 |
| Recording type * Age | F(2,96.74 = 0.61) | **0.021** |

In summary, the main effects reported by Attaheri et al. 2022 (1) were replicated in the second half of the sample, apart from some main effects of age being inconsistent when comparing the first and second halves of the cohort for the ∼4.05Hz power peak. Complete consistency regarding age was only achieved for the 4.35Hz power peak, which was the measure taken forward.

#### 3.1.2 Cortical tracking (mTRF) analyses

A linear mixed effects model (LMEM) was constructed using the same parameters as described in Attaheri et al., 2022 (1) to examine potential developmental changes in cortical tracking in the delta, theta and alpha (control) bands. The full sample submitted to the mTRF LMEM was as follows, 4-months (N=108), 7-months (N=104) and 11-months (N=106). Main effects of data type (real *r* values, chance *r* values), frequency band (delta, theta or alpha) and age (4-, 7- or 11-months) were investigated along with interactions between data type by frequency band, data type by age and age by frequency band. A random intercept (AR1) per participant was included to account for individual differences across the 3 recording sessions (4-, 7- and 11-months).

The main pattern of results replicated between the first and second halves of the sample, with significant fixed effects of data type and frequency band, as well as significant interactions between data type by frequency band and age by frequency band (Table 4). There was no significant effect of age when analysing the second half of the sample in isolation, although this effect was present in the first half of the sample (Table 4). When using the full cohort, tests of fixed effects showed significant effects of data type, frequency band and age. These results indicate that the real *r* values (cortical tracking) in response to nursery rhymes were significantly above randomly permuted *r* values and that the cortical tracking values differed by frequency band and age. The significant interaction of frequency band by data type indicated that cortical tracking in the delta and theta bands was driven by their values being significantly above the randomly permuted *r* values (as compared to the alpha base case; Table S3). The marginal means across the three ages, revealed that cortical tracking estimates (*r* values) were significantly higher in both delta (random permuted *r* values, *M* = 0.015, *SE* ± 0.001; real *r* values, *M* = 0.03, *SE* ± 0.001), and theta bands (random permuted *r* values, *M* = 0.011, *SE* ± 0.001; real *r* values, *M* = 0.016, *SE* ± 0.001), compared to the alpha base case (random permuted *r* values, *M* = 0.007, *SE* ± 0.001; real *r* values, *M* = 0.007, *SE* ± 0.001). Furthermore, alpha tracking was again not significantly above chance (Table S3) when using the full sample, mimicking the results from the first half of the sample (1). The significant interaction between age and frequency band showed that the relative differences in r values (both real and random) varied across the three age groups and frequency bands. In summary, the main cortical tracking effects reported by Attaheri et al. 2022 (1) were replicated in the second half of the sample.

**Table 4.**
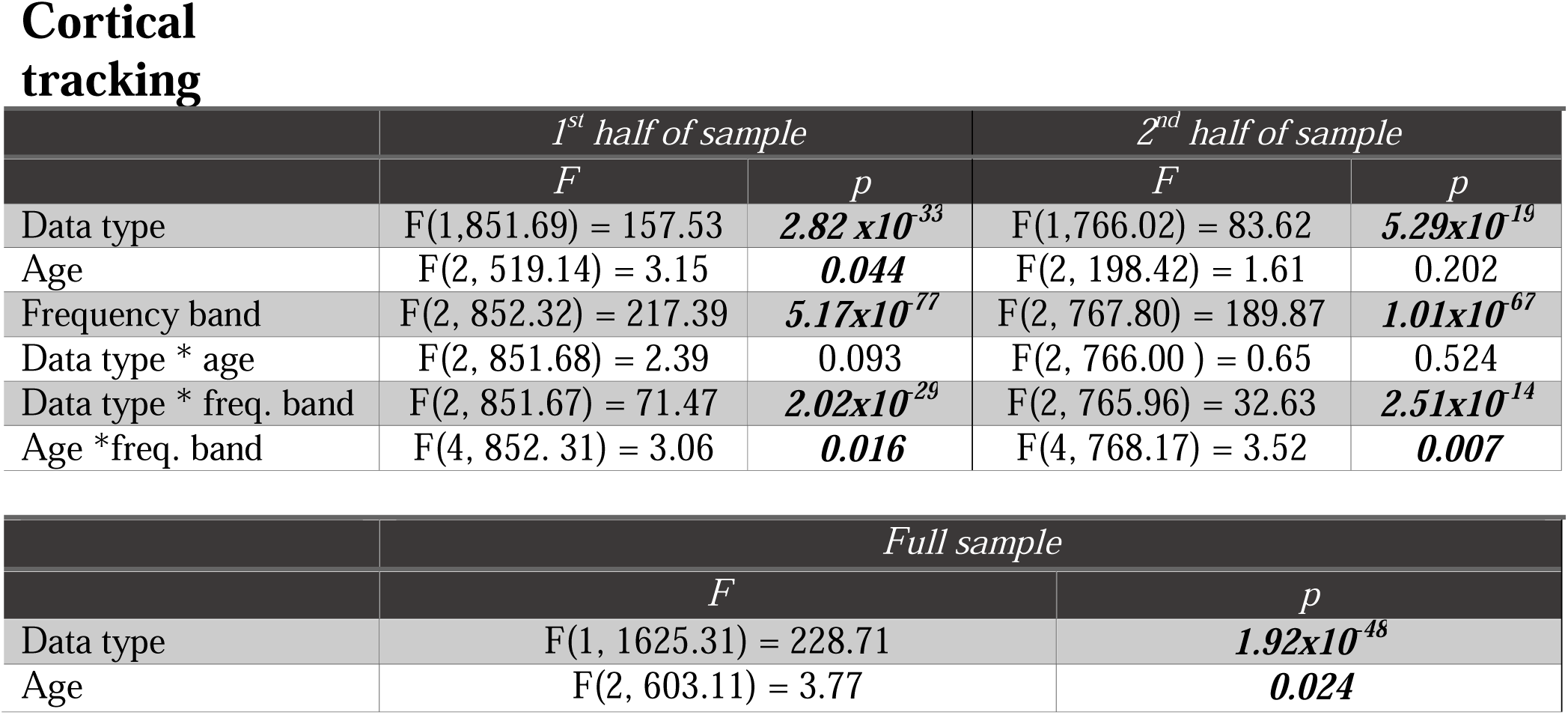
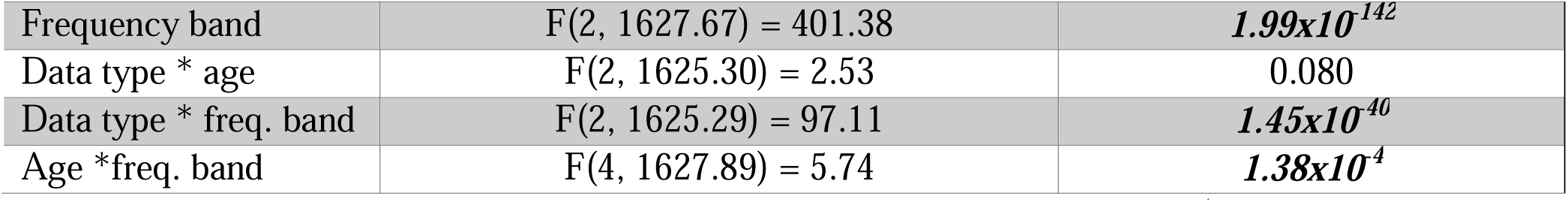
Tests of fixed effects for LMEM of cortical tracking across data type (real r values, random permutation *r* values), frequency band (delta, theta or alpha) and age (4-, 7- or 11- months). Results reported for separate LMEMs run on the first half, second half and full sample of data. Significant results (p<0.05) denoted by ***bold italic text***. The Satterthwaite approximation was applied to approximate the degrees of freedom, due to the missing cases.

Cortical tracking
|  | *1<sup>st</sup> half of sample* |  | *2<sup>nd</sup> half of sample* |  |
| --- | --- | --- | --- | --- |
|  | *F* | *p* | *F* | *p* |
| Data type | F(1,851.69) = 157.53 | **$2.82 \times 10^{-33}$** | F(1,766.02) = 83.62 | **$5.29 \times 10^{-19}$** |
| Age | F(2, 519.14) = 3.15 | **0.044** | F(2, 198.42) = 1.61 | 0.202 |
| Frequency band | F(2, 852.32) = 217.39 | **$5.17 \times 10^{-77}$** | F(2, 767.80) = 189.87 | **$1.01 \times 10^{-67}$** |
| Data type * age | F(2, 851.68) = 2.39 | 0.093 | F(2, 766.00) = 0.65 | 0.524 |
| Data type * freq. band | F(2, 851.67) = 71.47 | **$2.02 \times 10^{-29}$** | F(2, 765.96) = 32.63 | **$2.51 \times 10^{-14}$** |
| Age *freq. band | F(4, 852.31) = 3.06 | **0.016** | F(4, 768.17) = 3.52 | **0.007** |

Table 4. Tests of fixed effects for LMEM of cortical tracking across data type (real $r$ values, random permutation $r$ values), frequency band (delta, theta or alpha) and age (4-, 7- or 11-months).
|  | *Full sample* |  |
| --- | --- | --- |
|  | *F* | *p* |
| Data type | F(1, 1625.31) = 228.71 | **$1.92 \times 10^{-48}$** |
| Age | F(2, 603.11) = 3.77 | **0.024** |
| Frequency band | F(2, 1627.67) = 401.38 | ***1.99x10<sup>-142</sup>*** |
| Data type * age | F(2, 1625.30) = 2.53 | 0.080 |
| Data type * freq. band | F(2, 1625.29) = 97.11 | ***1.45x10<sup>-40</sup>*** |
| Age *freq. band | F(4, 1627.89) = 5.74 | ***1.38x10<sup>-4</sup>*** |

Taken together, the replication of PSD and cortical tracking results in the lower frequencies strengthens the conclusion that delta and theta EEG responses track the acoustic envelope of sung and chanted speech from the age of 4-months and that this infant neural response is robust and reliable.

#### 3.1.3 Phase amplitude coupling (PAC) analysis

The PAC analyses (Fig. 3) were conducted for the second half of the sample and the full cohort, replicating the LMEM conducted with the first half of the sample (1). The full sample submitted to the PAC LMEM was as follows, 4-months (N=108), 7-months (N=103) and 11- months (N=107). This LMEM included fixed effect factors of data type (real nMI values or chance nMI values), low-frequency phase (delta or theta), high-frequency amplitude (beta or gamma) and age (4-, 7- or 11-months). Interactions of data type by age, by low-frequency phase, data type by high-frequency amplitude, data type by age, low-frequency phase by age, high-frequency amplitude by age and low-frequency phase by high-frequency amplitude were also included in the model, with a random intercept (AR1) per participant (see methods section and Attaheri et al., 2022 (1) for detailed methods). Tests of fixed effects are reported in Table 5 and estimates of fixed effects are provided in Table S4. The LMEM intended to explore whether the nMI values were significantly above chance and if there were any significant differences when either delta or theta was the low-frequency carrier phase or when either beta or gamma were the high-frequency amplitudes, and if this relationship was affected by age.

**Figure 2.**
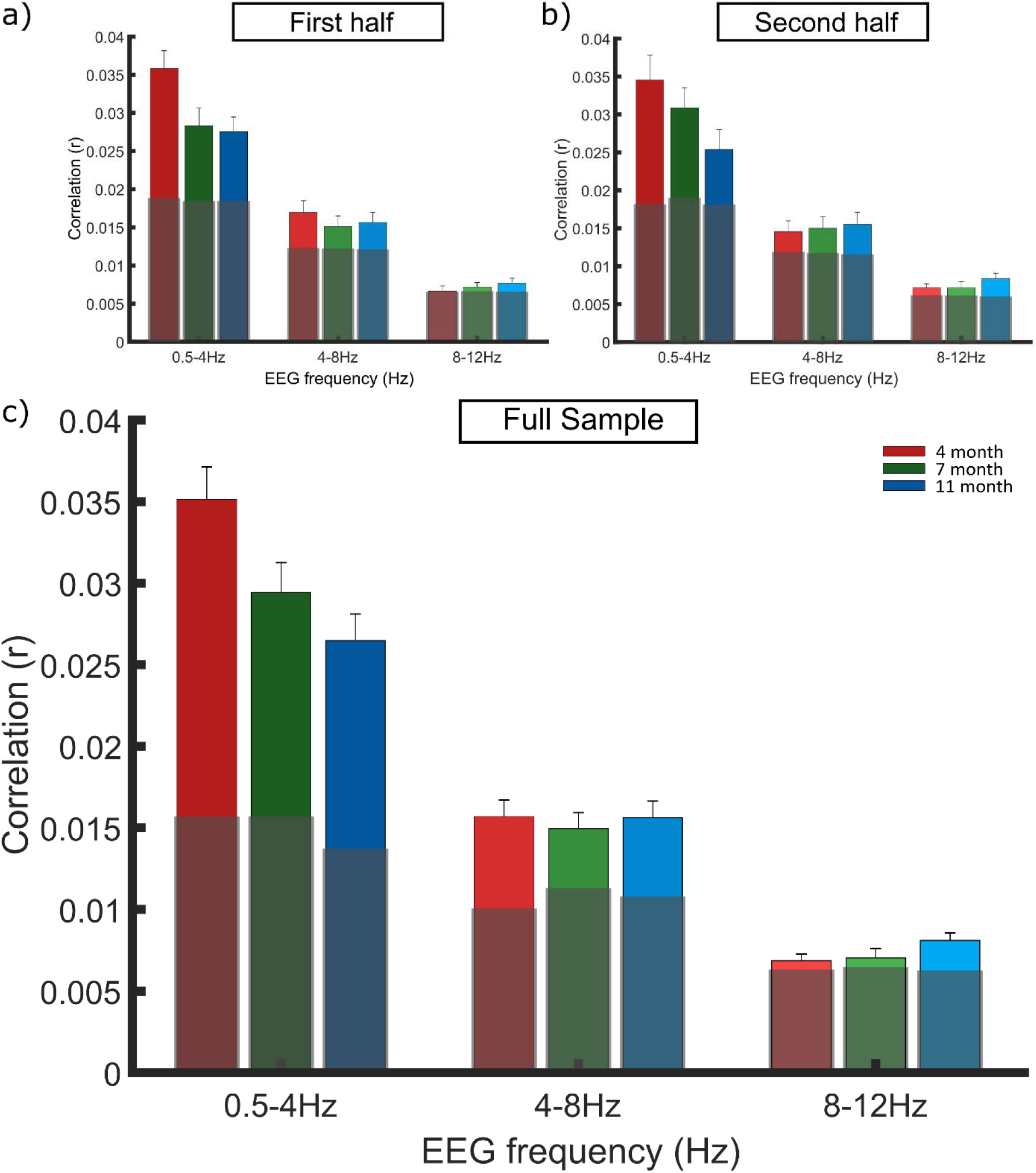
mTRF replication. Grand averaged stimulus reconstruction values (Pearson’s r) per frequency band in the a) first half, b) second half and c) full infant cohort. Each bar represents the grand average r value across the participants at the 3 age points: 4-months (red), 7-months (green) and 11-months (blue). Age responses are grouped along the X axis by EEG frequency band, delta (0.5–4 Hz) theta (4–8 Hz) and alpha (8–12 Hz). Light grey boxes signify the average chance *r* value across subjects, calculated separately for each band. Outlier analysis was conducted to remove extreme values which would compromise the LMEMs. Panel a is reproduced from Attaheri et al. (2022), with permission.

**Figure 3.**
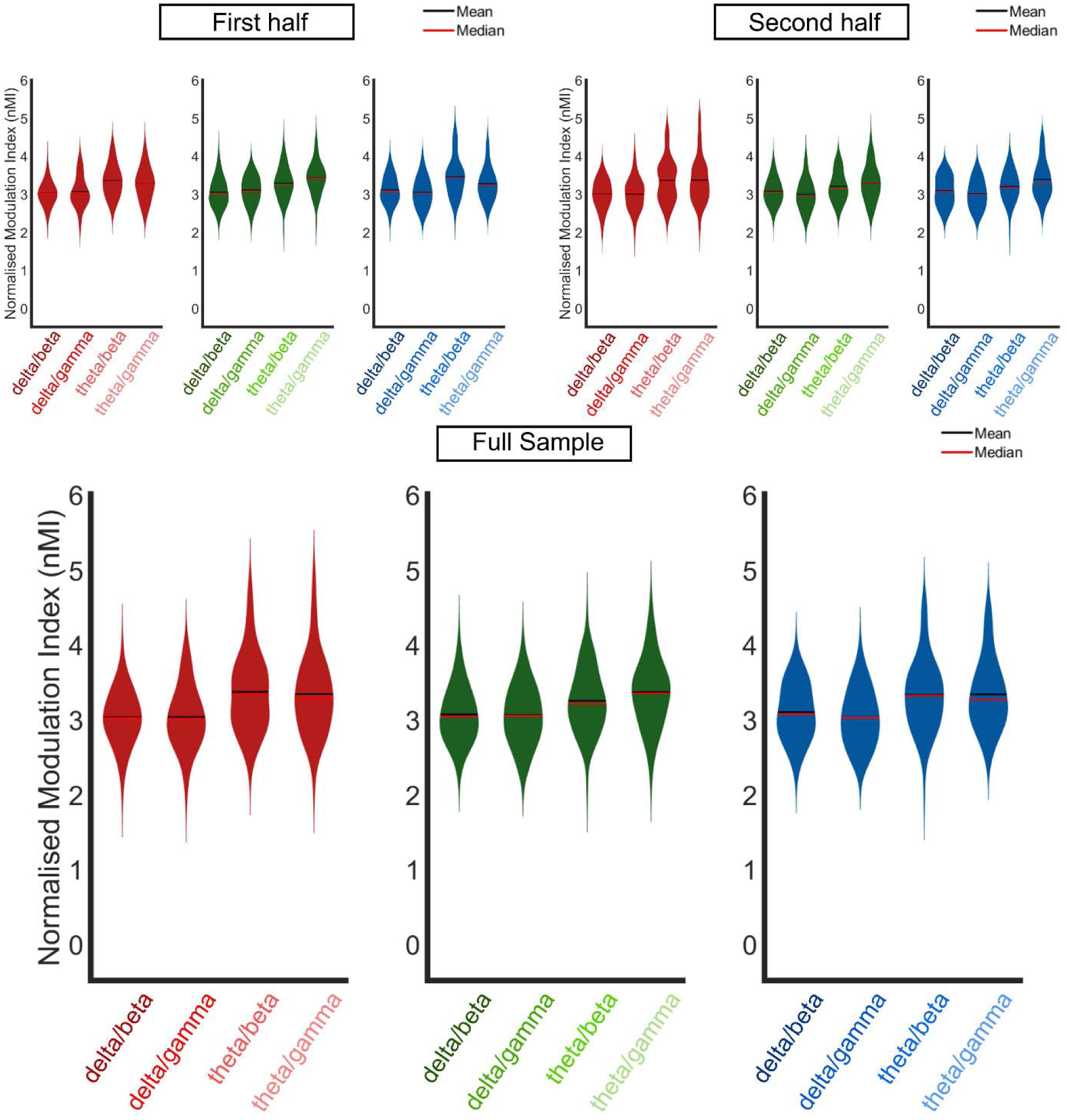
PAC replication. Violin plots of phase amplitude coupling (normalised modulation index, nMI) at 4-months (red), 7-months (green) and 11-months (blue) for the a) first half, b) second half and c) full infant cohort. The nMIs, from all significant analysis windows, were averaged for each individual infant. The PAC pairing with the maximum nMI per infant, from within the pre-defined frequency bands of interest (delta/beta, delta/gamma, theta/beta and theta/gamma; delta 2–4 Hz, theta 4–8 Hz, beta 15–30 Hz and gamma 30–45 Hz), were included in the grand average violin plot. Horizontal black and red lines represent the group mean and median respectively. Outlier analysis was conducted to remove extreme data points that would compromise the LMEM. Panel A is reproduced from Attaheri et al. 2022 (1), with permission.

**Table 5.** Tests of fixed effects generated from LMEM, assess how PAC (nMI) varied across data types (real or random nMI values), low-frequency phase (LFP), high-frequency amplitude (HFA) and age (4-, 7- or 11-months). Results reported for separate LMEMs run on the first half, second half and full sample of data. Significant results (p<0.05) denoted by **bold italic text**. The Satterthwaite approximation was applied to approximate the degrees of freedom, due to the missing cases.

|  | *1<sup>st</sup> half of sample* |  | *2<sup>nd</sup> half of sample* |  |
| --- | --- | --- | --- | --- |
|  | *F* | *p* | *F* | *p* |
| Data type (rand vs real nMI) | $F(1, 1178.08) = 4921.51$ | **<0.001** | $F(1, 1113.52) = 4409.87$ | **<0.001** |
| Low freq. phase (LFP) | $F(1, 1178.08) = 55.79$ | **<0.001** | $F(1, 1113.52) = 64.77$ | **<0.001** |
| Age | $F(2, 686.71) = .06$ | 0.941 | $F(2, 202.36) = 1.81$ | 0.167 |
| High freq. amp (HFA) | $F(1, 1178.08) = .05$ | 0.824 | $F(1, 1113.52) = 1.68$ | 0.196 |
| Data type * LFP | $F(1, 1178.08) = 54.34$ | **<0.001** | $F(1, 1113.52) = 62.86$ | **<0.001** |
| Data type * HFA | $F(1, 1178.08) = .00$ | 0.949 | $F(1, 1113.52) = 4.00$ | **0.046** |
| Data type * Age | $F(2, 1178.08) = 1.68$ | 0.187 | $F(2, 1113.52) = 3.26$ | **0.039** |
| LFP * Age | $F(2, 1178.08) = .47$ | 0.626 | $F(2, 1113.52) = 2.10$ | 0.123 |
| HFA * Age | $F(2, 1178.08) = 2.01$ | 0.134 | $F(2, 1113.52) = .28$ | 0.757 |
| LFP * HFA | $F(1, 1178.08) = .11$ | 0.738 | $F(1, 1113.52) = 2.48$ | 0.115 |
| Age * LFP * HFA | $F(2, 1178.08) = .64$ | 0.526 | $F(2, 1113.52) = .69$ | 0.504 |

|  | *Full sample* |  |
| --- | --- | --- |
|  | *F* | *p* |
| Data type (rand vs real nMI) | $F(1, 2269.31) = 9217.24$ | **<0.001** |
| Low freq. phase (LFP) | $F(1, 2269.31) = 118.88$ | **<0.001** |
| Age | $F(2, 790.87) = 1.40$ | 0.247 |
| High freq. amp (HFA) | $F(1, 2269.31) = 1.10$ | 0.294 |
| Data type * LFP | $F(1, 2269.31) = 115.42$ | **<0.001** |
| Data type * HFA | $F(1, 2269.31) = 1.78$ | 0.182 |
| Data type * Age | $F(2, 2269.31) = 2.31$ | 0.100 |
| LFP * Age | $F(2, 2269.31) = 2.21$ | 0.110 |
| HFA * Age | $F(2, 2269.31) = 1.79$ | 0.167 |
| LFP * HFA | F(1, 2269.31) = 0.71 | 0.399 |
| Age * LFP * HFA | F(1, 2269.31) = 0.44 | 0.642 |

The main pattern of results replicated between the first and second halves of the sample, with significant fixed effects of data type and low-frequency phase, as also reflected in the full cohort. The main effect of LFP arose because the nMI values were larger when theta (M =2.568, SE ± 0.013) was the low-frequency phase coupling with high-frequency amplitudes rather than delta (M = 2.405, SE ± 0.013, Table 5). A non-significant effect of HFA suggested that the beta and gamma amplitudes coupled equally as strongly with delta and theta phases (Table 5). A significant interaction between data type and LFP was also observed. Whilst both delta and theta displayed significant above-chance PAC, the significant interaction of LFP by data type reflected the fact that the relative difference between chance and real nMI values was significantly greater in the theta band than in the delta band, delta (chance nMI, M=1.770, SE ± 0.016; real nMI, M = 3.041, SE ± 0.016), Theta (chance nMI, M=1.772, SE ± 0.016; real nMI, M = 3.364, SE ± 0.016). Whilst the main pattern of results replicated between the first and second halves of the sample, one inconsistency was that the second half of the sample also showed a significant interaction between data type and HFA and between data type and age. However, as no main effects of age nor HFA were observed in the second half of the sample, these results are difficult to interpret.

#### 3.1.4 Theta/delta PSD power ratio

A significant difference between the theta/delta PSD power ratio across the three ages was also found (Fig. 4). Ratio values significantly increased with age (4 months, *M* = 0.546, *SE* ± 0.027 ; 7 months, *M* = 0.78, *SE* ± 0.073; 11 months, *M* = 1.017, *SE* ± 0.110), indicated by a one-way ANOVA (F(2,273) = 9.54, *p* = 9.9×10^-5^). The ratio increased because the amount of theta PSD power increased relative to the amount of delta PSD power as our sample aged from 4- to 11- months (see Figure 1). For comparison, the child modelling using EEG recorded from children with and without dyslexia aged around 9 years showed that a higher theta/delta ratio was associated with poorer phonological awareness (39). In the filtered speech intervention study with dyslexic children (34), in which delta-band modulations in natural speech were amplified via filtering, the theta/delta ratio improved when the children were listening to the filtered speech (compared to natural speech) because delta power increased. Accordingly, the theta/delta ratio appears to be a sensitive index of speech-related neural dynamics in infants and children.

**Figure 4.**
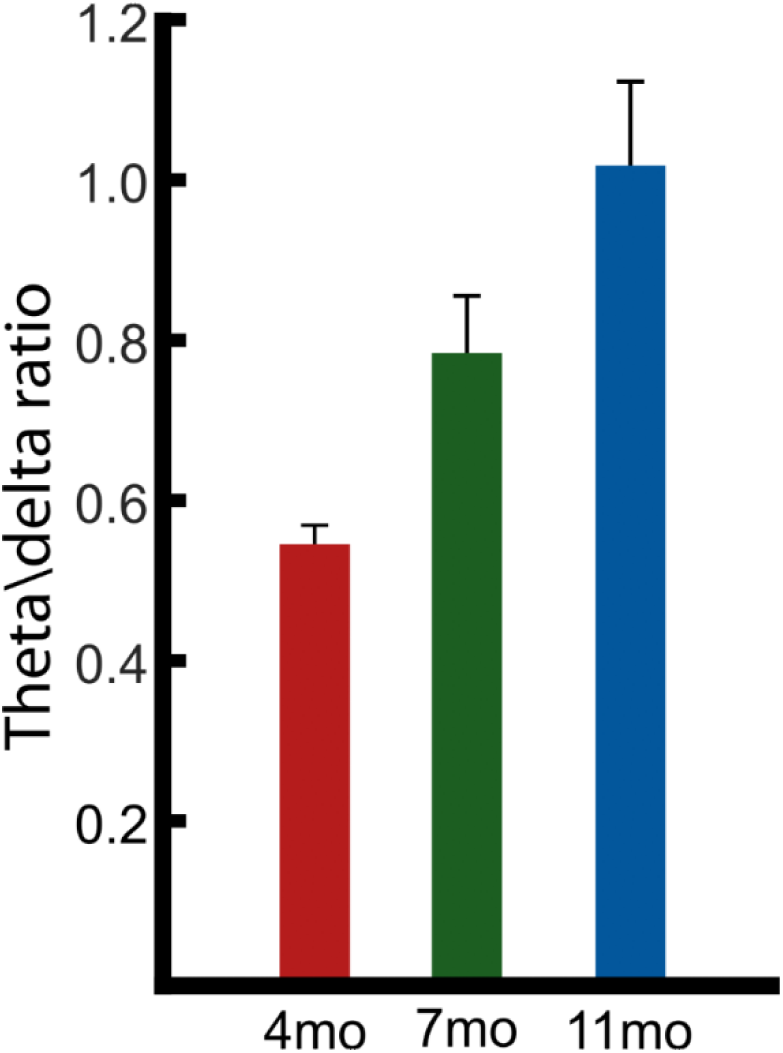
Theta / delta PSD ratio. The PSD peak per individual at 4.35 Hz and 1.92 Hz, taken from the spectral decomposition of the EEG signal, was used to create a theta / delta ratio calculation (4.35/1.92). Data provided at 4-months (red), 7-months (green) and 11-months (blue) with standard error bars in black.

### 3.2 Neural measures as predictors of language acquisition

Multivariate and univariate linear models were used to test whether the neural findings reported in 3.1 (for the full sample) were related to the different language measures using the analysis strategy described at 2.1 above. As explained in Rocha et al., (4), these language measures were selected as most reliable in previous analyses of the Cambridge UK BabyRhythm cohort, as they had the largest number of infants contributing data and scores were neither at floor nor ceiling (4). As noted, the parent-estimated measures were receptive and productive vocabulary scores from the UK-CDI at 24 months (49) and the infant-led measures were (i) The ability to point at 12- months, (ii) Word recognition at 18-months on a computerised infant-operated comprehension test (CCT), and (iii-v) Phonological processing at 24-months, as measured by non-word repetition of consonants, syllables and stress patterns (NWR). Due to the inherent difference in parent-estimated and infant-led measures of language, separate multivariate and univariate linear models were run for each set of scores. The results of these analyses for both parent-estimated and infant-led outcomes are reported in Tables 6-11. The univariate analyses often included more cases, as the multivariate analyses only included infants who contributed data points to all the language measures as well as to the neural measure being analysed.

**Table 6.** Multivariate and univariate linear models, investigating whether ∼1.92 Hz PSD power at 11 months predicted either infant-led or parent-estimated language outcomes. The table details the multivariate linear models followed by the univariate linear models describing whether ∼1.92 Hz PSD power at 11 months predicted, a) infant-led language measures or b) the parent-estimated language measures. Bonferroni correction for multiple comparisons in the univariate models led to modified significant alpha levels of, p = <0.01 for infant-led and p = <0.025 for parent-estimated (denoted by ***bold italic text***). Modified alpha level for trends are p = <0.02 for infant-led and p = <0.05 for parent-estimated (denoted by **bold text**).

a, Infant-led language measures
| Multivariate linear model | $Df_{num}$ | $Df_{denom}$ | $F$ | $p$ |
| --- | --- | --- | --- | --- |
| Infant-led language measures | 5 | 65 | 1.257 | 0.29 |
| Univariate linear models | $Df$ | $\beta_{estimate}$ | $SE$ | $p$ |
| NWR Consonants, 24 months | 1,79 | $5.23 \times 10^{-5}$ | $8.50 \times 10^{-5}$ | 0.95 |
| NWR Syllables, 24 months | 1,79 | $5.54 \times 10^{-4}$ | $8.56 \times 10^{-4}$ | 0.53 |
| NWR Stress, 24 months | 1,79 | $5.51 \times 10^{-4}$ | $7.83 \times 10^{-4}$ | 0.48 |
| CCT, 18 months | 1,79 | $7.16 \times 10^{-4}$ | $6.32 \times 10^{-4}$ | 0.26 |
| Pointing, 12 months | 1,90 | 0.003 | 0.001 | **0.014** |

b, Parent-estimated language measures
| Multivariate linear model | $Df_{num}$ | $Df_{denom}$ | $F$ | $p$ |
| --- | --- | --- | --- | --- |
| Parent-estimated language measures | 2 | 75 | 0.560 | 0.57 |
| Univariate linear models | $Df$ | $\beta_{estimate}$ | $SE$ | $p$ |
| CDI comprehension, 24 months | 1,76 | 0.373 | 0.445 | 0.40 |
| CDI production, 24 months | 1,76 | 0.179 | 0.465 | 0.70 |

**Table 7.** Multivariate and univariate linear models, investigating whether ∼4.35 Hz PSD power at 11 months predicted either infant-led or parent-estimated language outcomes. The table details the multivariate linear models followed by the univariate linear models describing whether ∼4.35 Hz PSD power at 11 months predicted a) infant-led language measures or b) the parent-estimated language measures. Bonferroni correction for multiple comparisons in the **univariate models** led to modified significant alpha levels of, p = <0.01 for infant-led and p = <0.025 for parent-estimated (denoted by ***bold italic text***). Modified alpha level for trends are p = <0.02 for infant-led and p = <0.05 for parent-estimated (denoted by **bold text**).

a, Infant-led language measures
| Multivariate linear model | $Df_{num}$ | $Df_{denom}$ | $F$ | $p$ |
| --- | --- | --- | --- | --- |
| Infant-led language measures | 5 | 65 | 1.76 | 0.13 |
| Univariate linear models | $Df$ | $\beta_{estimate}$ | $SE$ | $p$ |
| NWR Consonants, 24 months | 1,79 | $-4.93 \times 10^{-4}$ | $3.69 \times 10^{-4}$ | 0.19 |
| NWR Syllables, 24 months | 1,79 | $-8.36 \times 10^{-4}$ | $3.65 \times 10^{-4}$ | 0.025 |
| NWR Stress, 24 months | 1,79 | $-5.31 \times 10^{-4}$ | $3.40 \times 10^{-4}$ | 0.12 |
| CCT, 18 months | 1,79 | $-1.62 \times 10^{-4}$ | $2.76 \times 10^{-4}$ | 0.56 |
| Pointing, 12 months | 1,90 | $-2.95 \times 10^{-4}$ | $-5.98 \times 10^{-4}$ | 0.62 |

b, Parent-estimated language measures
| Multivariate linear model | $Df_{num}$ | $Df_{denom}$ | $F$ | $p$ |
| --- | --- | --- | --- | --- |
| Parent-estimated language measures | 2 | 75 | 3.18 | **0.047** |
| Univariate linear models | $Df$ | $\beta_{estimate}$ | $SE$ | $p$ |
| CDI comprehension, 24 months | 1,76 | -0.086 | 0.209 | 0.68 |
| CDI production, 24 months | 1,76 | -0.346 | 0.215 | 0.11 |

**Table 8.** Multivariate and univariate linear models, investigating whether delta/theta ratio of PSD power (∼1.92 Hz/4.35 Hz) at 11 months predicted either infant-led or parent-estimated language outcomes. The table details the multivariate linear models followed by the univariate linear models describing whether delta/theta ratio of PSD power (∼1.92 Hz/4.35 Hz) at 11 months predicted a) infant-led language measures or b) the parent-estimated language measures. Bonferroni correction for multiple comparisons in the **univariate models** led to modified significant alpha levels of, p = <0.01 for infant-led and p = <0.025 for parent-estimated (denoted by ***bold italic text***). Modified alpha level for trends are p = <0.02 for infant-led and p = <0.05 for parent-estimated (denoted by **bold text)**.

a, Infant-led language measures
| Multivariate linear model | $Df_{num}$ | $Df_{denom}$ | $F$ | $p$ |
| --- | --- | --- | --- | --- |
| Infant-led language measures | 5 | 65 | 2.68 | **0.029** |
| Univariate linear models | $Df$ | $\beta_{estimate}$ | $SE$ | $p$ |
| NWR Consonants, 24 months | 1,79 | -0.019 | 0.032 | 0.56 |
| NWR Syllables, 24 months | 1,79 | -0.058 | 0.032 | 0.07 |
| NWR Stress, 24 months | 1,79 | -0.044 | 0.029 | 0.14 |
| CCT, 18 months | 1,79 | -0.023 | 0.023 | 0.32 |
| Pointing, 12 months | 1,90 | -0.065 | 0.049 | 0.19 |

b, Parent-estimated language measures
| Multivariate linear model | $Df_{num}$ | $Df_{denom}$ | $F$ | $p$ |
| --- | --- | --- | --- | --- |
| Parent-estimated language measures | 2 | 75 | 3.65 | **0.031** |
| Univariate linear models | $Df$ | $\beta_{estimate}$ | $SE$ | $P$ |
| CDI comprehension, 24 months | 1,76 | -6.760 | 16.44 | 0.68 |
| CDI production, 24 months | 1,76 | -28.55 | 16.82 | 0.094 |

**Table 9.**
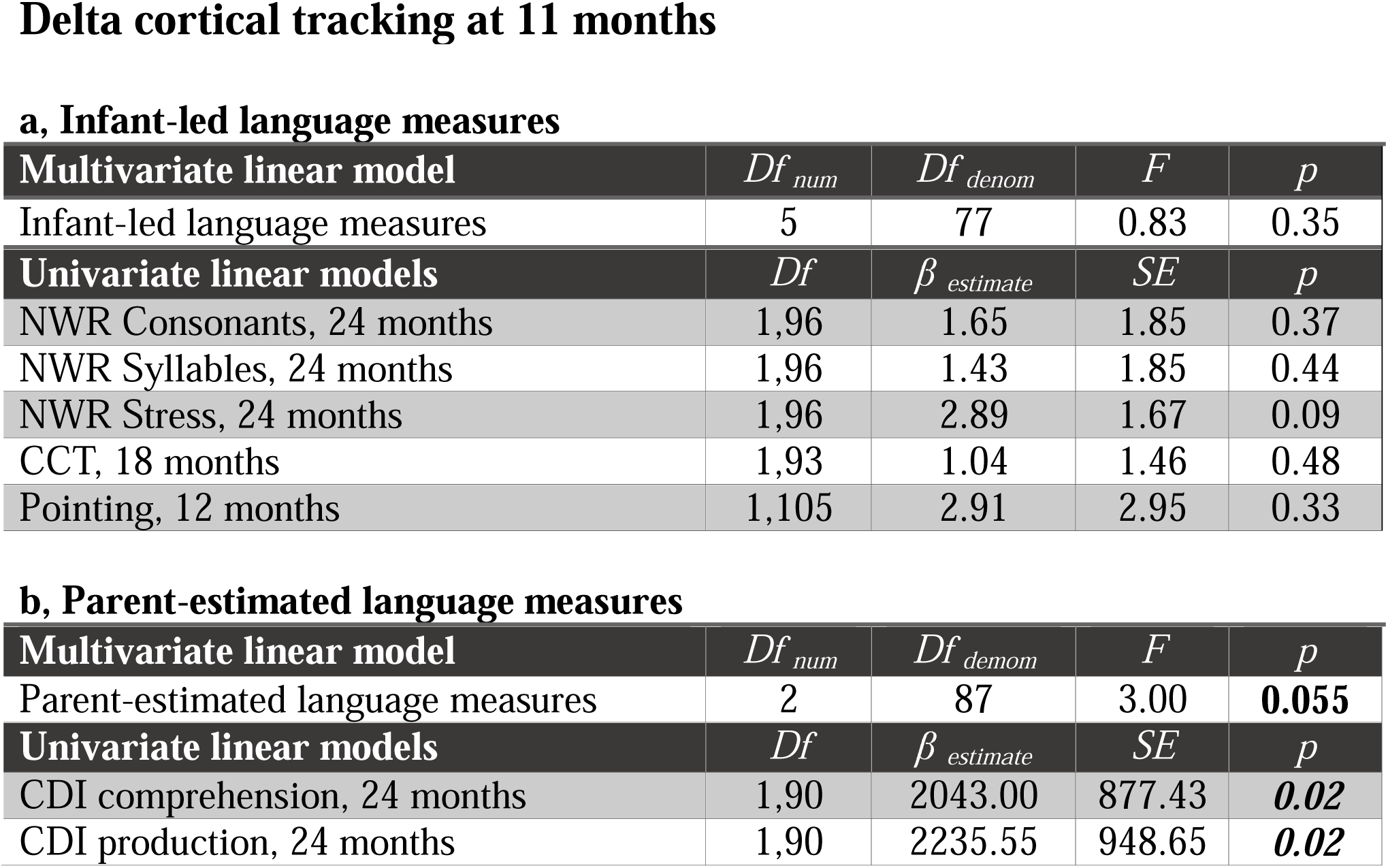
Multivariate and univariate linear models, investigating whether delta cortical tracking at 11 months predicted either infant-led or parent-estimated language outcomes. The table details the multivariate linear models followed by the univariate linear models describing whether delta cortical tracking at 11 months predicted a) infant-led language measures or b) the parent-estimated language measures. Bonferroni correction for multiple comparisons in the **univariate models** led to modified significant alpha levels of, p = <0.01 for infant-led and p = <0.025 for parent-estimated (denoted by ***bold italic text***). Modified alpha level for trends are p = <0.02 for infant-led and p = <0.05 for parent-estimated (denoted by **bold text**).

**Table 10.** Multivariate and univariate linear models, investigating whether theta cortical tracking at 11 months predicted either infant-led or parent-estimated language outcomes. The table details the multivariate linear models followed by the univariate linear models describing whether theta cortical tracking at 11 months predicted a) infant-led language measures or b) the parent-estimated language measures. Bonferroni correction for multiple comparisons in the **univariate models** led to modified significant alpha levels of, p = <0.01 for infant-led and p = <0.025 for parent-estimated (denoted by ***bold italic text***). Modified alpha level for trends are p = <0.02 for infant-led and p = <0.05 for parent-estimated (denoted by **bold text**).

a, Infant-led language measures
| Multivariate linear model | $Df_{num}$ | $Df_{demom}$ | $F$ | $p$ |
| --- | --- | --- | --- | --- |
| Infant-led language measures | 5 | 77 | 0.89 | 0.50 |
| Univariate linear models | $Df$ | $\beta_{estimate}$ | $SE$ | $p$ |
| NWR Consonants, 24 months | 1,96 | -4.32 | 2.91 | 0.14 |
| NWR Syllables, 24 months | 1,96 | -3.51 | 2.92 | 0.23 |
| NWR Stress, 24 months | 1,96 | -2.24 | 2.67 | 0.40 |
| CCT, 18 months | 1,93 | -2.72 | 2.28 | 0.24 |
| Pointing, 12 months | 1,105 | 5.72 | 4.61 | 0.22 |

b, Parent-estimated language measures
| Multivariate linear model | $Df_{num}$ | $Df_{demom}$ | $F$ | $p$ |
| --- | --- | --- | --- | --- |
| Parent-estimated language measures | 2 | 87 | 0.25 | 0.78 |
| Univariate linear models | $Df$ | $\beta_{estimate}$ | $SE$ | $p$ |
| CDI comprehension, 24 months | 1,90 | -828.31 | 1452.9 | 0.57 |
| CDI production, 24 months | 1,90 | -1114.12 | 1570.3 | 0.48 |

**Table 11.** Multivariate and univariate linear models, investigating whether delta-beta, delta-gamma, theta-beta or theta-gamma PAC at 4 months predicted either infant-led or parent-estimated language outcomes. The table details the multivariate linear models followed by the univariate linear models describing whether PAC at 4 months predicted a) infant-led language measures or b) the parent-estimated language measures. Bonferroni correction for multiple comparisons in the **univariate models** led to modified significant alpha levels of, p = <0.01 for infant-led and p = <0.025 for parent-estimated (denoted by ***bold italic text***). Modified alpha level for trends are p = <0.02 for infant-led and p = <0.05 for parent-estimated (denoted by **bold text**).

a, Infant-led language
|  | *Delta-beta* |  | *Delta-gamma* |  | *Theta-beta* |  | *Theta-gamma* |  |
| --- | --- | --- | --- | --- | --- | --- | --- | --- |
| **Multivariate linear model** | *F* | *p* | *F* | *p* | *F* | *p* | *F* | *p* |
| Infant-led language measures | 0.81 | 0.54 | 0.45 | 0.81 | 0.90 | 0.49 | 4.13 | **0.002** |
|  | *Delta-beta* |  | *Delta-gamma* |  | *Theta-beta* |  | *Theta-gamma* |  |
| **Univariate linear models** | $\beta_{est.}$ | *P* | $\beta_{est.}$ | *P* | $\beta_{est.}$ | *p* | $\beta_{est.}$ | *p* |
| NWR Consonants, 24 months | -0.11 | 0.17 | 0.05 | 0.51 | -0.04 | 0.52 | 0.08 | 0.09 |
| NWR Syllables, 24 months | -0.09 | 0.27 | 0.03 | 0.70 | -0.04 | 0.49 | 0.14 | **0.002** |
| NWR Stress, 24 months | -0.07 | 0.37 | 0.02 | 0.79 | -0.08 | 0.05 | 0.0. | 0.65 |
| CCT, 18 months | -0.13 | 0.03 | 0.03 | 0.54 | 0.04 | 0.39 | 0.06 | 0.05 |
| Pointing, 12 months | -0.1 | 0.41 | -0.03 | 0.78 | -0.08 | 0.39 | 0.02 | 0.77 |

b, Parent-estimated language
|  | *Delta-beta* |  | *Delta-gamma* |  | *Theta-beta* |  | *Theta-gamma* |  |
| --- | --- | --- | --- | --- | --- | --- | --- | --- |
| **Multivariate linear model** | *F* | *p* | *F* | *p* | *F* | *p* | *F* | *p* |
| Parent-est. language measures | 3.38 | **.039** | 2.68 | **0.07** | 1.46 | 0.24 | 4.86 | **0.01** |
|  | *Delta-beta* |  | *Delta-gamma* |  | *Theta-beta* |  | *Theta-gamma* |  |
| **Univariate linear models** | $\beta_{est.}$ | *p* | $\beta_{est.}$ | *p* | $\beta_{est.}$ | *p* | $\beta_{est.}$ | *p* |
| CDI comprehension, 24 mo | -53.4 | 0.20 | 85.0 | **0.02** | -26.5 | 0.38 | 24.2 | 0.28 |
| CDI production, 24 mo | -103.3 | **0.02** | 77.1 | 0.06 | 3.35 | 0.92 | 60.1 | **0.01** |

#### 3.2.1. PSD and language outcomes

The 11-months EEG data were selected for comparison to both the parent-estimated and infant-led language measures, due to the observed developmental maturation of the ∼4 Hz peak (see Figure 1). As described above, the 4.35 Hz peak was the more robust and consistent of the two theta peaks (see Tables 1 and 2). It was therefore used for predicting the language outcomes. The analyses for the 4-month and 7-month EEG are also provided as Tables S5 to S11 (with 4.05 Hz peak results reported in Tables S7-S9). The analyses for the ∼1.92 Hz peak are also reported here. 77 infants provided complete data for all the parent-estimated measures and PSD power results, whilst 70 infants provided data for all the infant-led language measures and PSD power.

Regarding global language outcomes, the multivariate models indicated that increased ∼1.92 Hz PSD power in response to nursery rhymes did not lead to significant prediction of either infant-led (F(5,65) = 1.257, *p=* 0.285) nor parent-estimated (F(2,75) = 0.560, *p=* 0.574) outcomes (Table 6). However, the univariate linear models revealed a trend for pointing at 12-months to be correlated with increases in ∼1.92 Hz power (Table 6a; *p=* 0.014, Bonferroni corrected trend α values, *p* = <0.02 for infant-led univariate models).

The multivariate analyses for ∼4.35 Hz PSD power in response to nursery rhymes showed no effect on language outcomes for the infant-led measures (F(5,65) = 1.758, *p=* 0.134, Table 7a). This was also the case in the univariate analyses. However, for the parent-estimated measures, increased theta power led to a significant *decrease* (F(2,75) = 3.18, *p=* 0.047) in the global parent-estimated language outcomes (Table 7b). It is important to note that whilst the univariate linear model analyses indicated that comprehension and production did not individually make significant contributions to this prediction (Table 7b), the direction of the beta estimates suggest that higher power was associated with fewer known words later in development. Accordingly there is a global effect of theta power on parent-estimated (UK-CDI) language outcomes, but this is not driven by either single parent-estimated measure.

A further pair of multivariate linear model analyses showed that a larger theta/delta PSD power ratio (4.35 Hz / 1.92 Hz) led to a decrease in performance for both infant-led (F(5,65) = 2.684, *p=* 0.029, Table 8a) and parent-estimated (F(2,75) = 3.65, *p=* 0.031, Table 8b) global outcomes. The more theta power that was present relative to delta power, the worse the infants’ performance on either type of outcome. Again, the univariate linear models suggested that these global effects were not driven by any single individual measure. However, it was measures of language *production* that showed *p* values <.10 (NWR of syllables, *p*= .07, CDI production, *p*= .09).

#### 3.2.2 mTRF and language outcomes

The 11-months EEG data were selected for analyses with both the parent-estimated and infant-led language measures, as these were recorded the closest in time to the language measures (please see Tables S12-15 for matching data for the earlier age groupings; these did not reach significance). 89 infants provided complete data for all the parent-estimated measures and a cortical tracking value, whilst 82 infants provided data for all the infant-led language measures and a cortical tracking value.

The multivariate linear models regarding the parent-estimated language measures (Table 9a) displayed a trend (not significant at an alpha value of 0.05) towards better language outcomes, with increased accuracy of delta-band cortical tracking in response to nursery rhymes predicting better outcomes (F(2,87) = 3.00, *p=* 0.055). No multivariate effect was present for the infant-led measures, (F(5,77) = 0.83, *p=* 0.350). However, the univariate linear model analyses showed that both CDI comprehension and CDI production at 24 months were significantly predicted by delta band cortical tracking (*p*’s= 0.02, Table 9b). By contrast, the accuracy of theta band cortical tracking did not predict either infant-led (F(5,77) = 0.886, *p=* 0.495) or parent-estimated (F(2,87) = 0.250, *p=* 0.780) language outcomes (Table 10). This was unexpected given the adult literature. As alpha cortical oscillations did not show above chance cortical tracking, comparison to language measures were not considered (however the data are provided in Tables S16-S18 for completeness)

#### 3.2.3 PAC and language outcomes

Phase amplitude coupling (PAC) at 4-months was selected for the language outcome analyses as PAC did not show any developmental changes indicating that the early presence of stronger PAC may be informative (please see Tables S19-20 for matching data for the later age groupings). 87 infants provided complete data for all the parent-estimated measures and a PAC value, whilst 81 infants provided data for all the infant-led language measures and a PAC value. Multivariate linear models were run separately to investigate whether delta-beta, delta-gamma, theta-beta or theta-gamma PAC respectively predicted the language outcome measures.

Considering delta as the LFP first, increased delta-beta PAC predicted a global *decrease* in the parent-estimated (F(2,85) = 3.38, *p=* 0.039) language measures (Table 11b). The univariate linear models indicated that CDI production at 24 months drove this overall effect (Table 11b). Increased delta-beta PAC did not predict the infant-led language outcome measures in the multivariate analysis (F(5,76) = 0.447, *p=* 0.815) nor in the univariate analyses (Table 11a). By contrast, stronger delta-gamma PAC showed a trend to increase parent-estimated language measures (F(2,85) = 2.68, *p=* 0.075), with the significant univariate linear model indicating that CDI comprehension at 24 months drove this trend (*p=0.02).* Delta-gamma PAC did not predict the infant-led language outcome measures (F(5,76) = 0.447, *p=* 0.815, Table 11a), and the delta-gamma univariate models showed similar results.

When theta was the LFP, the multivariate linear models indicated that increased theta-gamma PAC predicted a significant global increase in the infant-led language measures (F(5,76) = 4.126, *p=* 0.002, Table 11a) and the parent-estimated (F(2,85) = 4.86, *p=* 0.010) language measures (Table 11b). Bonferroni-corrected univariate linear models highlighted that better non-word repetition of syllables at 24-months (*p=* .002) was the strongest contributing factor. Theta-beta PAC showed no effect on later language outcomes in the multivariate analyses, both for the infant-led measures (F(5,76) = 0.897, *p=* 0.488) and the parent-estimated measures (F(2,85) = 146, *p=* 0.240, Table 11b). The univariate analyses showed similar results.

In summary, theta-gamma PAC was associated with a significant increase in both the infant-led and parent-estimated language outcomes, in both the multivariate and univariate analyses. Trend effects suggested that delta-gamma PAC also contributed to increases in parent-estimated language outcomes. Conversely, delta-beta PAC predicted a significant global *decrease* in parent-estimated language measures, and theta-beta PAC exerted no effects. The data suggest that the coupling of LFPs to gamma-band HFAs could be a key factor in language acquisition.

## 4 Discussion

According to recent extensions of Temporal Sampling theory (13,37) cortical tracking of infant directed speech (IDS) by delta- and theta-band neural signals and delta-theta neural dynamics may be key factors in explaining individual differences in the developmental trajectories for language acquisition. In the Cambridge UK BabyRhythm infant sample assessed here (N=113), individual differences in delta-band cortical tracking (in the univariate analyses), theta-gamma PAC (in both univariate and multivariate analyses), theta power (in the multivariate analyses) and theta/delta power ratio (multivariate analyses) during the first year of life were all significant predictors of language outcomes in infants’ second year of life. These infant data are consistent with neural oscillatory factors that have been found to explain individual differences in language development in children, for example the fidelity and synchronicity of delta-band encoding of speech envelope information for phonological development (35,60). They are also consistent with a recent EEG modelling study based on child recordings during passive speech listening, which showed that a higher theta/delta power ratio was related to phonological difficulties in dyslexia (39), and a child MEG study which showed that amplifying speech information in the delta band reduced this power ratio when dyslexic children were listening to filtered speech (34). Further, the infant neural oscillatory factors identified here were broadly consistent in terms of replicating our previous findings with the first half of the infant cohort (1). The presence and maturation of low-frequency (<12 Hz) cortical oscillations (PSD), cortical speech tracking (mTRF) and phase amplitude coupling (PAC) in response to sung IDS was largely similar across both halves of the longitudinal Cambridge UK BabyRhythm cohort.

The replication of stronger delta band tracking (Figure 2) than theta band tracking in the youngest (4-month-old) infants suggests that delta band cortical tracking may be a key early building block for language acquisition (21). Efficient delta-band cortical tracking would enable infant brains to lock on to the information-rich strong (or stressed) syllables in the speech signal, acoustic landmarks that occur on average every 500ms (2Hz) across languages (61) . This mechanism would enable the infant brain to begin to represent the acoustic structure of human speech (59,62). Dynamic interactions with theta-band cortical tracking may then support further linguistic learning (see below). The importance of delta-band cortical tracking at 11 months for language outcomes in our univariate analyses (Table 9) is consistent with child MEG studies, which show that typically-developing children exhibit efficient delta-band tracking of speech-in-noise, but not efficient theta-band tracking (63). The finding that delta band cortical tracking at 11 months predicted early language comprehension and production in the univariate analyses (Table 9) is also congruent with research in the adult brain, where delta-band tracking is associated with key language-related skills such as perceptual grouping (64,65), discourse-level parsing related to phrasing (66,67) and comprehending speech-in-noise (68).

In many adult studies, the theta band is considered the key linguistic oscillator (6). In our infant cohort, better theta-gamma PAC at 4-months was associated with better outcomes in both the infant-led and parent-estimated multivariate analyses (assessing global language outcomes) and the univariate analyses (Table 11). Adult work shows how PAC could operate as a neural mechanism for grouping fine-grained phonetic information into syllabic units (22,26,29). The univariate analyses for theta-gamma PAC revealed that nonword repetition, a phonological output measure (specifically syllable production at 24-months) and CDI production (parental estimation of vocabulary output) were strong drivers of these effects (Table 11). By contrast, delta-based PAC predicted increases in the language outcome measures when gamma was the HFA (parent-estimated univariate analyses), but *decreases* in the language outcome measures when beta was the HFA (parent-estimated multivariate and univariate analyses). Further, not all the effects found for theta-band cortical tracking were positive in nature. Theta-band PSD increased significantly during the first year of life, and greater theta-band PSD at 11 months was associated with poorer language outcomes as measured by parental estimates of comprehension and production at 24-months in the multivariate analyses (Table 7). A larger theta/delta PSD power ratio at 11 months was also associated with poorer language outcomes (Table 8), as measured by multivariate analyses of both parent-estimated and infant-led measures. This latter finding underscores the importance of considering neural dynamics when assessing developmental trajectories. In the previous EEG modelling work with 9-year-old children with and without dyslexia that investigated theta/delta ratios, a dyslexia-specific pattern of delta and theta power dynamics during natural speech processing was seen to become progressively more atypical the worse the child’s phonological awareness, and this was not related to age (32). Further, the theta/delta ratio could be improved during story listening for children with dyslexia by increasing delta power via a filtered speech manipulation (34).

The infant literature is sparse regarding PAC dynamics. For adults, theta-gamma PAC has been suggested as a neural mechanism for processing phonetic information in the speech signal (6,69,70). When the same rhythmic speech stimuli used in the current study were presented to adult participants, theta/gamma coupling was stronger than delta/gamma coupling (28). For our infant participants presented with the same nursery rhyme stimuli, theta-gamma PAC at 4 months was the consistent predictor of language outcomes in both the multivariate and the univariate analyses (Table 11). Indeed, it is the phase of theta band rather than delta band oscillations that adjusts to speech rate variations for adults, coupling with gamma amplitudes (26). Whilst both delta and theta phases couple with higher-frequency amplitudes in the infant brain, it seems that theta-based PAC is more strongly predictive of early language outcomes. The univariate analyses (Table 11) suggested that measures involving speech production, namely nonword repetition and productive vocabulary, were important regarding these relations. The adult PAC literature suggests that delta-beta PAC is particularly associated with measures of speech production via audio-motor dynamics (27,71). In our data, stronger delta-beta PAC before 1 year of age was associated with poorer language outcomes at 24 months (Table 11). In the univariate analyses, the negative association was for CDI production (*p*= .02). This may be a transitory developmental effect. As the infants in our cohort continue to develop and produce more speech, it may be that delta-beta PAC will begin to play a positive role in predicting language outcomes.

The current study has several limitations. Recording EEG with infants is an inherently noisy process, and this affected our data, as different infants had to be excluded from different analyses, reducing sample size for the longitudinal language outcome assessments, particularly regarding the multivariate analyses. A key source of noisy data are infant fussing and movement, and noise was particularly apparent in the 7-month-old recordings for our sample. This could be age-related, as by 7 months of age infants have a much larger range of movement. A second limitation is that even though our study followed a relatively large cohort of infants (122 recruited and 113 retained), a study with a larger participant pool may have had increased chances of uncovering developmentally important time-lagged associations. Finally, the grand averaged peaks observed at 4.05 Hz and 4.35Hz may correspond to a developmental maturation of the PSD peak between 7-and 11-month-olds. The current study adopted a data-driven approach across conditions and ages, however, future studies could include a separate condition to identify endogenous peaks for each age group. This information could then be used to define the analysis peaks in the experimental condition. Notwithstanding the limitations of the study, here we present a large longitudinal sample demonstrating that individual differences in neural dynamics in response to speech stimuli predict some aspects of early infant language acquisition.

## 5 Conclusion

Estimates of the efficiency of low-frequency cortical tracking, PSD, phase-amplitude coupling and theta-delta power dynamics in an infant’s first year of life predict their early language outcomes. Importantly, the neural mechanisms measured here are automatic processes that are inherent to the human brain, and as such may represent physiological priors for language learning in a Bayesian sense (72). Accordingly, individual differences in the efficiency of these mechanisms in the pre-verbal stage of language learning assessed here are unlikely to reflect top-down linguistic processes such as language comprehension (18). Delta-band cortical tracking at 11 months in the univariate analyses and theta−gamma PAC at 4 months of age in both the univariate and multivariate analyses significantly predicted language comprehension and production at 24 months, with the latter predicting both word production and nonword repetition in the univariate analyses. Conversely, increased theta PSD power and a larger theta/delta ratio were associated with decreased performance on the language outcome measures in our multivariate analyses. While other studies have also indicated a predictive relationship between cortical tracking efficiency in infancy and language development (30), the current study is the first to show that individual differences in PAC and theta/delta power dynamics also predict infant language outcomes.

## Author Contributions

*Adam Attaheri*: EEG Paradigm Development, EEG Preprocessing, Investigation - Data Curation, Formal Analysis - design, creation and implementation of analysis, Writing - Original Draft.

*Áine Ní Choisdealbha*: EEG Preprocessing - Investigation - Data Curation - Writing - Review and Editing

*Sinead Rocha:* Investigation - Data Curation, Writing - Review and Editing

*Perrine Brusini*: EEG Paradigm Development, Investigation - Data Curation

*Giovanni Di Liberto*: Formal analysis, Writing - Review and Editing

*Natasha Mead*: Investigation - Data Curation

*Helen Olawole-Scott*: Investigation - Data Curation

*Panagiotis Boutris*: Investigation - Data Curation

*Samuel Gibbon*: Investigation - Data Curation

*Isabel Williams*: Investigation - Data Curation

*Christina Grey*: Investigation - Data Curation

*Maria Alfaro e Oliveira -* Data Curation

*Carmel Brough -* Data Curation Shelia Flanagan – Data curation

*Usha Goswami*: Conceptualization - Methodology, Funding Acquisition, Supervision, Project Administration, Writing - Original Draft

## Data Sharing agreement

The analyses were conducted by using publicly available MATLAB toolboxes that can be downloaded at http://audition.ens.fr/adc/NoiseTools/, https://sccn.ucsd.edu/eeglab/download.php, https://github.com/mickcrosse/mTRF-Toolbox.

The custom code used for the mTRF, PSD and PAC analysis can be downloaded from https://osf.io/ea9bd/. The final data included in the manuscript figures and statistics can also be downloaded from https://osf.io/ea9bd/.

## Acknowledgements

We would like to thank all the families and infants who kindly donated their time to this project. We would also like to thank Henna Ahmed for their time spent as an RA on the Cambridge UK BabyRhythm project. Finally, we would like to acknowledge Mike X Cohen’s statistical textbooks and online materials for guiding our statistical analysis.

## Supporting information captions

Figure S1. Grand average spectral decomposition of the EEG signal (1-12 Hz) of the full sample.

Figure S2. Grand average spectral decomposition of the EEG signal (1-12 Hz) of the second half of the sample.

Figure S3. Grand average spectral decomposition of the EEG signal (1-12 Hz) of the first half of the sample.

Figure S4, Average modulation spectrum of all the Nursery rhyme stimuli.

Figure S5, Example Modulation spectrum of one the Nursery rhyme stimuli (‘Simple Simon’).

Figure S6, Grand average of the 18 individual nursery rhymes “All Band AVG” Modulation spectrums.

Figure S7, Spectral decomposition of the EEG signal averaged across the full infant cohort (1–12 Hz plotted, from 0.5 to 45 Hz calculated), in response to the silent state period.

Table S1. Mean and Standard error of the PSD peak amplitudes, for both recording Silent state and Nursery rhyme recording conditions.

Table S2. Parameter estimates from the full sample PSD LMEM.

Table S3. Parameter estimates from the full sample cortical tracking LMEM.

Table S4. Parameter estimates from the full sample PAC LMEM.

Table S5. Multivariate and univariate linear models for 1.92 Hz PSD power at 4 months.

Table S6. Multivariate and univariate linear models for ∼1.92 Hz PSD power at 7 months.

Table S7. Multivariate and univariate linear models for ∼4.05 Hz PSD power at 4 months.

Table S8. Multivariate and univariate linear models for ∼4.05 Hz PSD power at 7 months.

Table S9. Multivariate and univariate linear models for ∼4.05 Hz PSD power at 11 months.

Table S10. Multivariate and univariate linear models for ∼4.35 Hz PSD power at 4 months.

Table S11. Multivariate and univariate linear models for ∼4.35 Hz PSD power at 7 months.

Table S12. Multivariate and univariate linear models for delta cortical tracking at 4 months.

Table S13. Multivariate and univariate linear models for delta cortical tracking at 7 months.

Table S14. Multivariate and univariate linear models for theta cortical tracking at 4 months.

Table S15. Multivariate and univariate linear models for theta cortical tracking at 7 months.

Table S16. Multivariate and univariate linear models for alpha cortical tracking at 4 months.

Table S17. Multivariate and univariate linear models for alpha cortical tracking at 7 months.

Table S18. Multivariate and univariate linear models for alpha cortical tracking at 11 months.

Table S19. Multivariate and univariate linear models for PAC at 7 months.

Table S20. Multivariate and univariate linear models for PAC at 11 months.

## References

1. Attaheri A, Choisdealbha ÁN, Di Liberto GM, Rocha S, Brusini P, Mead N, et al. Delta- and theta-band cortical tracking and phase-amplitude coupling to sung speech by infants. NeuroImage. 2022 Feb 15;118698.

2. Kuhl PK. Early language acquisition: cracking the speech code. Nat Rev Neurosci. 2004 Nov;5(11):831–43.

3. Mehler J, Jusczyk P, Lambertz G, Halsted N, Bertoncini J, Amiel-Tison C. A precursor of language acquisition in young infants. Cognition. 1988;29(2):143–78.

4. Rocha S, Attaheri A, Ní Choisdealbha Á, Brusini P, Mead N, Olawole-Scott H, et al. Precursors to infant sensorimotor synchronization to speech and non-speech rhythms: A longitudinal study. Dev Sci. 2024 Mar 12;n/a(n/a):e13483.

5. Obleser J, Kayser C. Neural Entrainment and Attentional Selection in the Listening Brain. Trends Cogn Sci. 2019 Nov 1;23(11):913–26.

6. Giraud AL, Poeppel D. Cortical oscillations and speech processing: emerging computational principles and operations. Nat Neurosci. 2012 Mar 18;15(4):511–7.

7. Ding N, Simon JZ. Cortical entrainment to continuous speech: functional roles and interpretations. Front Hum Neurosci [Internet]. 2014 [cited 2020 Jul 3];8. Available from: https://www.frontiersin.org/articles/10.3389/fnhum.2014.00311/full

8. Di Liberto GM, O’Sullivan JA, Lalor EC. Low-Frequency Cortical Entrainment to Speech Reflects Phoneme-Level Processing. Curr Biol CB. 2015 Oct 5;25(19):2457–65.

9. Luo H, Poeppel D. Phase patterns of neuronal responses reliably discriminate speech in human auditory cortex. Neuron. 2007 Jun 21;54(6):1001–10.

10. O’Sullivan JA, Power AJ, Mesgarani N, Rajaram S, Foxe JJ, Shinn-Cunningham BG, et al. Attentional Selection in a Cocktail Party Environment Can Be Decoded from Single-Trial EEG. Cereb Cortex N Y NY. 2015 Jul;25(7):1697–706.

11. Zion Golumbic EM, Ding N, Bickel S, Lakatos P, Schevon CA, McKhann GM, et al. Mechanisms Underlying Selective Neuronal Tracking of Attended Speech at a ‘Cocktail Party.’ Neuron. 2013 Mar 6;77(5):980–91.

12. Peelle J, Davis M. Neural Oscillations Carry Speech Rhythm through to Comprehension. Front Psychol. 2012;3:320.

13. Goswami U. Speech rhythm and language acquisition: an amplitude modulation phase hierarchy perspective. Ann N Y Acad Sci. 2019;1453(1):67–78.

14. Goswami U. A Neural Basis for Phonological Awareness? An Oscillatory Temporal-Sampling Perspective. Curr Dir Psychol Sci. 2018 Feb 1;27(1):56–63.

15. Baltzell LS, Srinivasan R, Richards VM. The effect of prior knowledge and intelligibility on the cortical entrainment response to speech. J Neurophysiol. 2017 Sep 6;118(6):3144–51.

16. Di Liberto GM, Crosse MJ, Lalor EC. Cortical Measures of Phoneme-Level Speech Encoding Correlate with the Perceived Clarity of Natural Speech. eNeuro [Internet]. 2018 Apr 16 [cited 2020 Jul 3];5(2). Available from: https://www.ncbi.nlm.nih.gov/pmc/articles/PMC5900464/

17. Millman RE, Johnson SR, Prendergast G. The Role of Phase-locking to the Temporal Envelope of Speech in Auditory Perception and Speech Intelligibility. J Cogn Neurosci. 2015 Mar 1;27(3):533–45.

18. Peelle JE, Gross J, Davis MH. Phase-locked responses to speech in human auditory cortex are enhanced during comprehension. Cereb Cortex N Y N 1991. 2013 Jun;23(6):1378–87.

19. Jessen S, Fiedler L, Münte TF, Obleser J. Quantifying the individual auditory and visual brain response in 7-month-old infants watching a brief cartoon movie. NeuroImage. 2019 Nov 15;202:116060.

20. Kalashnikova M, Peter V, Di Liberto GM, Lalor EC, Burnham D. Infant-directed speech facilitates seven-month-old infants’ cortical tracking of speech. Sci Rep [Internet]. 2018 Sep 13 [cited 2020 Jul 3];8. Available from: https://www.ncbi.nlm.nih.gov/pmc/articles/PMC6137049/

21. Ortiz Barajas MC, Guevara R, Gervain J. The origins and development of speech envelope tracking during the first months of life. Dev Cogn Neurosci. 2021 Jan 20;48:100915– 100915.

22. Gross J, Hoogenboom N, Thut G, Schyns P, Panzeri S, Belin P, et al. Speech Rhythms and Multiplexed Oscillatory Sensory Coding in the Human Brain. PLOS Biol. 2013 Dec 31;11(12):e1001752.

23. Canolty RT, Edwards E, Dalal SS, Soltani M, Nagarajan SS, Kirsch HE, et al. High Gamma Power Is Phase-Locked to Theta Oscillations in Human Neocortex. Science. 2006 Sep 15;313(5793):1626–8.

24. Canolty RT, Knight RT. The functional role of cross-frequency coupling. Trends Cogn Sci. 2010 Nov;14(11):506–15.

25. Lakatos P, Shah AS, Knuth KH, Ulbert I, Karmos G, Schroeder CE. An oscillatory hierarchy controlling neuronal excitability and stimulus processing in the auditory cortex. J Neurophysiol. 2005 Sep;94(3):1904–11.

26. Lizarazu M, Lallier M, Molinaro N. Phase−amplitude coupling between theta and gamma oscillations adapts to speech rate. Ann N Y Acad Sci. 2019 Oct;1453(1):140–52.

27. Arnal LH, Doelling KB, Poeppel D. Delta-Beta Coupled Oscillations Underlie Temporal Prediction Accuracy. Cereb Cortex N Y N 1991. 2014/05/20 ed. 2015 Sep;25(9):3077–85.

28. Attaheri A, Panayiotou D, Phillips A, Ní Choisdealbha Á, Di Liberto GM, Rocha S, et al. Cortical Tracking of Sung Speech in Adults vs Infants: A Developmental Analysis. Front Neurosci [Internet]. 2022;16. Available from: https://www.frontiersin.org/article/10.3389/fnins.2022.842447

29. Hyafil A, Fontolan L, Kabdebon C, Gutkin B, Giraud AL. Speech encoding by coupled cortical theta and gamma oscillations. Brownell H, editor. eLife. 2015 May 29;4:e06213.

30. Menn KH, Ward EK, Braukmann R, van den Boomen C, Buitelaar J, Hunnius S, et al. Neural Tracking in Infancy Predicts Language Development in Children With and Without Family History of Autism. Neurobiol Lang. 2022 Aug 17;3(3):495–514.

31. Kalashnikova M, Goswami U, Burnham D. Sensitivity to amplitude envelope rise time in infancy and vocabulary development at 3 years: A significant relationship. Dev Sci. 2019 Nov 1;22(6):e12836.

32. Araújo J, Simons BD, Varghese P, Mandke K, Kalashnikova M, Macfarlane A, et al. Atypical low-frequency cortical encoding of speech identifies children with developmental dyslexia. Front Hum Neurosci [Internet]. 2024;18. Available from: https://www.frontiersin.org/journals/human-neuroscience/articles/10.3389/fnhum.2024.1403677

33. Keshavarzi M, Di Liberto G, Gabrielczyk F, Wilson A, Macfarlane A, Goswami U. Atypical speech production of multisyllabic words by children with developmental dyslexia. 2022.

34. Mandke K, Flanagan S, Macfarlane A, Feltham G, Gabrielczyk F, Wilson AM, et al. Neural responses to natural and enhanced speech edges in children with and without dyslexia. Front Hum Neurosci [Internet]. 2023;17. Available from: https://www.frontiersin.org/articles/10.3389/fnhum.2023.1200950

35. Power AJ, Colling LJ, Mead N, Barnes L, Goswami U. Neural encoding of the speech envelope by children with developmental dyslexia. Brain Lang. 2016 Sep;160:1–10.

36. Power AJ, Mead N, Barnes L, Goswami U. Neural entrainment to rhythmic speech in children with developmental dyslexia. Front Hum Neurosci [Internet]. 2013 [cited 2020 Jul 3];7. Available from: https://www.frontiersin.org/articles/10.3389/fnhum.2013.00777/full

37. Goswami U. Language acquisition and speech rhythm patterns: an auditory neuroscience perspective. R Soc Open Sci. 2022 Jul 27;9.

38. Menn KH, Michel C, Meyer L, Hoehl S, Männel C. Natural infant-directed speech facilitates neural tracking of prosody. NeuroImage. 2022 May 1;251:118991.

39. Araújo J, Simons BD, Varghese P, Mandke K, Kalashnikova M, Macfarlane A, et al. Atypical encoding of speech identifies children with Dyslexia versus Developmental Language Disorder [Internet]. BioRxiv; Preprint. Available from: https://www.biorxiv.org/content/10.1101/2022.10.26.513864v1

40. Homae F, Watanabe H, Nakano T, Taga G. Large-Scale Brain Networks Underlying Language Acquisition in Early Infancy. Front Psychol [Internet]. 2011;2. Available from: https://www.frontiersin.org/articles/10.3389/fpsyg.2011.00093

41. Zoefel B, Oever ST, Sack AT. The Involvement of Endogenous Neural Oscillations in the Processing of Rhythmic Input: More Than a Regular Repetition of Evoked Neural Responses. 2018 Jun 6 [cited 2020 Jul 6]; Available from: https://www.repository.cam.ac.uk/handle/1810/276669

42. Mariani B, Nicoletti G, Barzon G, Ortiz Barajas MC, Shukla M, Guevara R, et al. Prenatal experience with language shapes the brain. Sci Adv [Internet]. 2023 Nov 22 [cited 2024 Sep 16];9(47). Available from: 10.1126/sciadv.adj3524

43. Ortiz-Barajas MC, Guevara R, Gervain J. Neural oscillations and speech processing at birth. iScience. 2023 Nov 17;26(11):108187.

44. Delorme A, Makeig S. EEGLAB: an open source toolbox for analysis of single-trial EEG dynamics including independent component analysis. J Neurosci Methods. 2004 Mar 15;134(1):9–21.

45. Kothe CA, Makeig S. BCILAB: a platform for brain–computer interface development. J Neural Eng. 2013 Aug 28;10(5):056014.

46. Crosse MJ, Di Liberto GM, Bednar A, Lalor EC. The Multivariate Temporal Response Function (mTRF) Toolbox: A MATLAB Toolbox for Relating Neural Signals to Continuous Stimuli. Front Hum Neurosci [Internet]. 2016 [cited 2020 Jul 3];10. Available from: https://www.frontiersin.org/articles/10.3389/fnhum.2016.00604/full

47. Pasley BN, David SV, Mesgarani N, Flinker A, Shamma SA, Crone NE, et al. Reconstructing Speech from Human Auditory Cortex. PLOS Biol. 2012 Jan 31;10(1):e1001251.

48. Özkurt TE, Schnitzler A. A critical note on the definition of phase–amplitude cross-frequency coupling. J Neurosci Methods. 2011 Oct 15;201(2):438–43.

49. Alcock K, Meints K, Rowland CF, Brelsford V, Christopher A, Just J. The UK Communicative Development Inventories: Words and Gestures - Manual and Norms. Emsworth, UK: J & R Press; 2020.

50. Liszkowski U, Carpenter M, Tomasello M. Reference and attitude in infant pointing. J Child Lang. 2007/01/25 ed. 2007;34(1):1–20.

51. Friend M, Keplinger M. An Infant-Based Assessment of Early Lexicon Acquisition. Behav Res Methods Instrum Comput J Psychon Soc Inc. 2003 Jun 1;35:302–9.

52. Hoff E, Core C, Bridges K. Non-word repetition assesses phonological memory and is related to vocabulary development in 20- to 24-month-olds. J Child Lang. 2008/10/06 ed. 2008;35(4):903–16.

53. Ní Choisdealbha Á, Attaheri A, Rocha S, Mead N, Olawole-Scott H, Brusini P, et al. Neural phase angle from two months when tracking speech and non-speech rhythm linked to language performance from 12 to 24 months. Brain Lang. 2023 Aug 1;243:105301.

54. Ní Choisdealbha Á, Attaheri A, Rocha S, Mead N, Olawole-Scott H, Alfaro e Oliveira M, et al. Cortical tracking of visual rhythmic speech by 5- and 8-month-old infants: Individual differences in phase angle relate to language outcomes up to 2 years. Dev Sci. 2024 Mar 14:e13502.

55. Marshall PJ, Bar-Haim Y, Fox NA. Development of the EEG from 5 months to 4 years of age. Clin Neurophysiol. 2002;113(8):1199–208.

56. Orekhova EV, Stroganova TA, Posikera IN, Elam M. EEG theta rhythm in infants and preschool children. Clin Neurophysiol Off J Int Fed Clin Neurophysiol. 2006 May;117(5):1047–62.

57. Stroganova TA, V. Orekhova E, Posikera IN. Externally and internally controlled attention in infants: an EEG study. Int J Psychophysiol. 1998 Nov 1;30(3):339–51.

58. Cuevas K, Bell MA. 293C13EEG Frequency Development Across Infancy and Childhood. In: Gable PA, Miller MW, Bernat EM, editors. The Oxford Handbook of EEG Frequency [Internet]. Oxford University Press; 2022 [cited 2023 Feb 26]. p. 0. Available from: 10.1093/oxfordhb/9780192898340.013.13

59. Leong V, Kalashnikova M, Burnham D, Goswami U. The Temporal Modulation Structure of Infant-Directed Speech. 2017 Mar 27 [cited 2020 Jul 3]; Available from: https://www.repository.cam.ac.uk/handle/1810/263724

60. Molinaro N, Lizarazu M, Lallier M, Bourguignon M, Carreiras M. Out-of-synchrony speech entrainment in developmental dyslexia. Hum Brain Mapp. 2016;37(8):2767–83.

61. Dauer RM. Stress-timing and syllable-timing reanalyzed**A preliminary version of this paper was read at the Annual Meeting of the Linguistic Society of America, San Antonio, Texas, December 28–30, 1980. The experimental work for this study was carried out at the Phonetics Laboratory, Edinburgh University. J Phon. 1983 Jan 1;11(1):51–62.

62. Leong V, Goswami U. Acoustic-Emergent Phonology in the Amplitude Envelope of Child-Directed Speech. PLOS ONE. 2015 Dec 7;10(12):e0144411.

63. Ghinst MV, Bourguignon M, Niesen M, Wens V, Hassid S, Choufani G, et al. Cortical Tracking of Speech-in-Noise Develops from Childhood to Adulthood. J Neurosci. 2019 Apr 10;39(15):2938–50.

64. Ding N, Melloni L, Zhang H, Tian X, Poeppel D. Cortical tracking of hierarchical linguistic structures in connected speech. Nat Neurosci. 2016 Jan;19(1):158–64.

65. Kösem A, van Wassenhove V. Distinct contributions of low- and high-frequency neural oscillations to speech comprehension. Lang Cogn Neurosci. 2017 May 28;32(5):536–44.

66. Boucher VJ, Gilbert AC, Jemel B. The role of low-frequency neural oscillations in speech processing: Revisiting delta entrainment. J Cogn Neurosci. 2019;31(8):1205–15.

67. Doelling KB, Arnal LH, Ghitza O, Poeppel D. Acoustic landmarks drive delta-theta oscillations to enable speech comprehension by facilitating perceptual parsing. NeuroImage. 2014 Jan 15;85 Pt 2:761–8.

68. Vanthornhout J, Decruy L, Wouters J, Simon JZ, Francart T. Speech Intelligibility Predicted from Neural Entrainment of the Speech Envelope. J Assoc Res Otolaryngol. 2018 Apr 1;19(2):181–91.

69. Fries P. The Model- and the Data-Gamma. Neuron. 2009 Dec 10;64(5):601–2.

70. Schroeder CE, Lakatos P. Low-frequency neuronal oscillations as instruments of sensory selection. Trends Neurosci. 2009 Jan 1;32(1):9–18.

71. Poeppel D, Assaneo MF. Speech rhythms and their neural foundations. Nat Rev Neurosci. 2020 Jun 1;21(6):322–34.

72. Doelling KB, Arnal LH, Assaneo MF. Adaptive oscillators provide a hard-coded Bayesian mechanism for rhythmic inference. bioRxiv. 2022 Jan 1;2022.06.18.496664.

