## Supplement for "Infant low-frequency EEG cortical power, cortical tracking and phase-amplitude coupling predicts language a year later"

### *PSD peak identification*

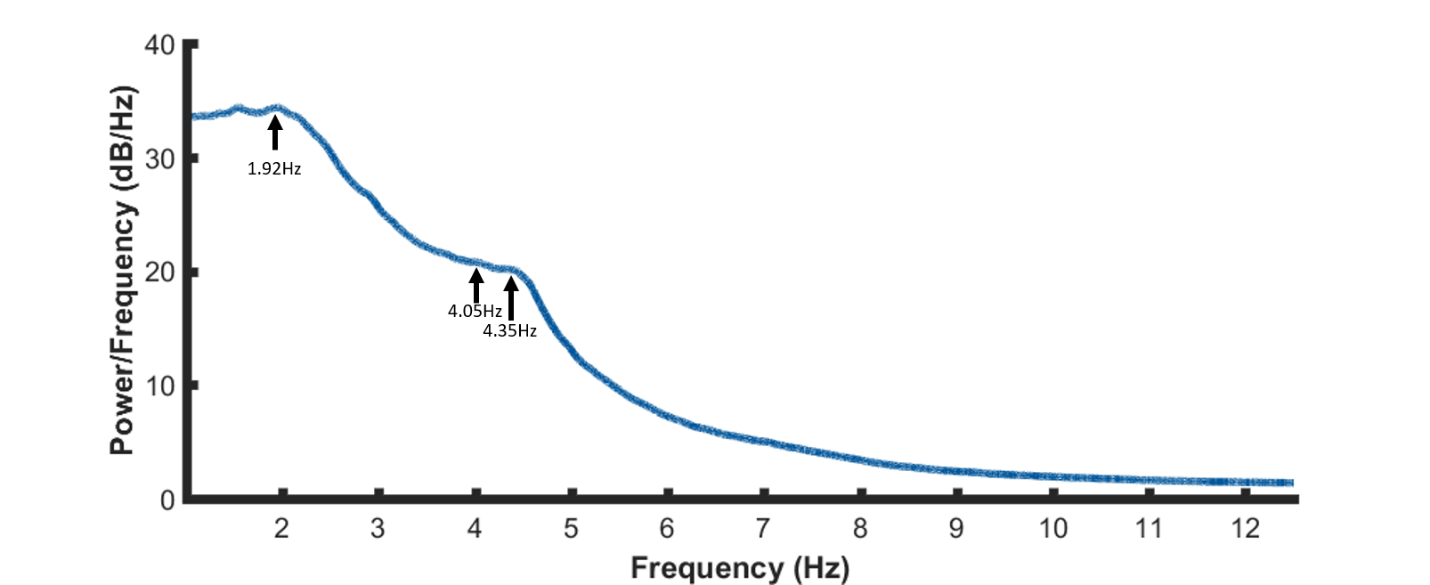

**Figure S1. Grand average spectral decomposition of the EEG signal (1-12 Hz) of the full sample.** A periodogram was used to obtain a power spectral density (PSD) estimate separately for 4-, 7- and 11-months data, before being averaged together to create a grand average. Arrows show the center frequency of peaks of interest used in main analysis. These were defined by visual inspection.

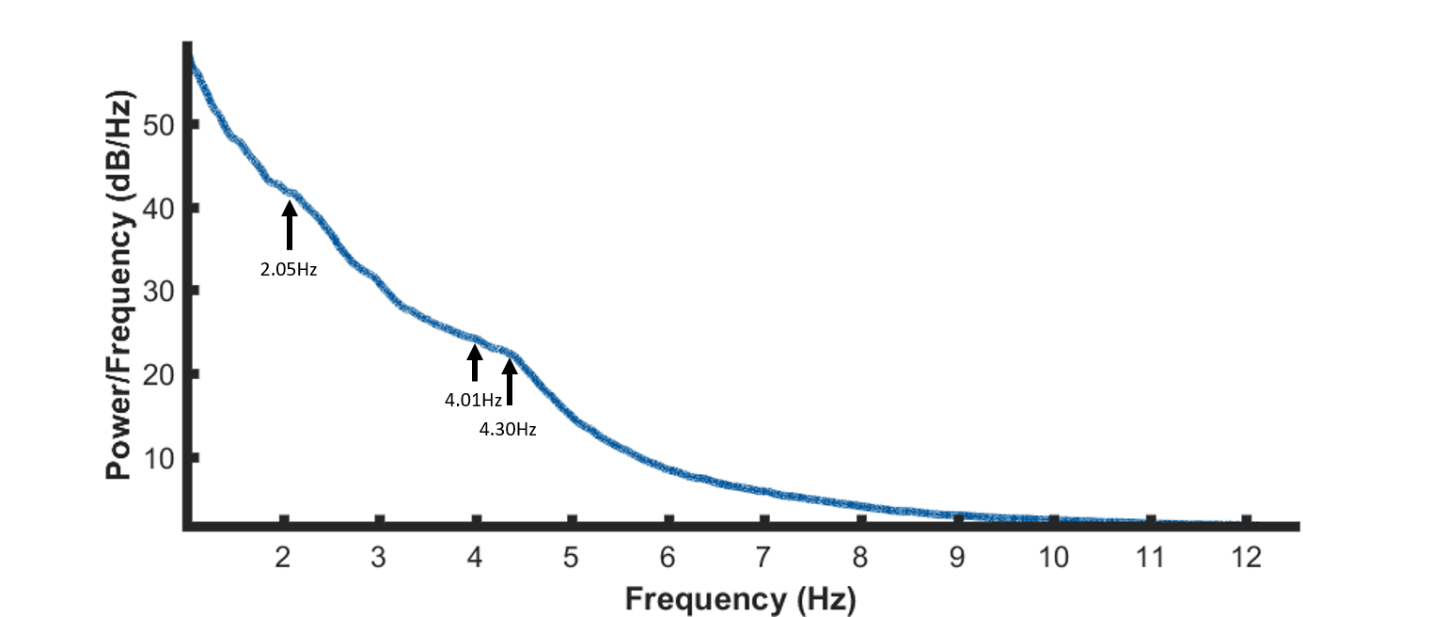

**Figure S2. Grand average spectral decomposition of the EEG signal (1-12 Hz) of the second half of the sample**. A periodogram was used to obtain a power spectral density (PSD) estimate separately for 4-, 7- and 11-months data, before being averaged together to create a grand average. Arrows show the center frequency of peaks of interest. These were defined by visual inspection.

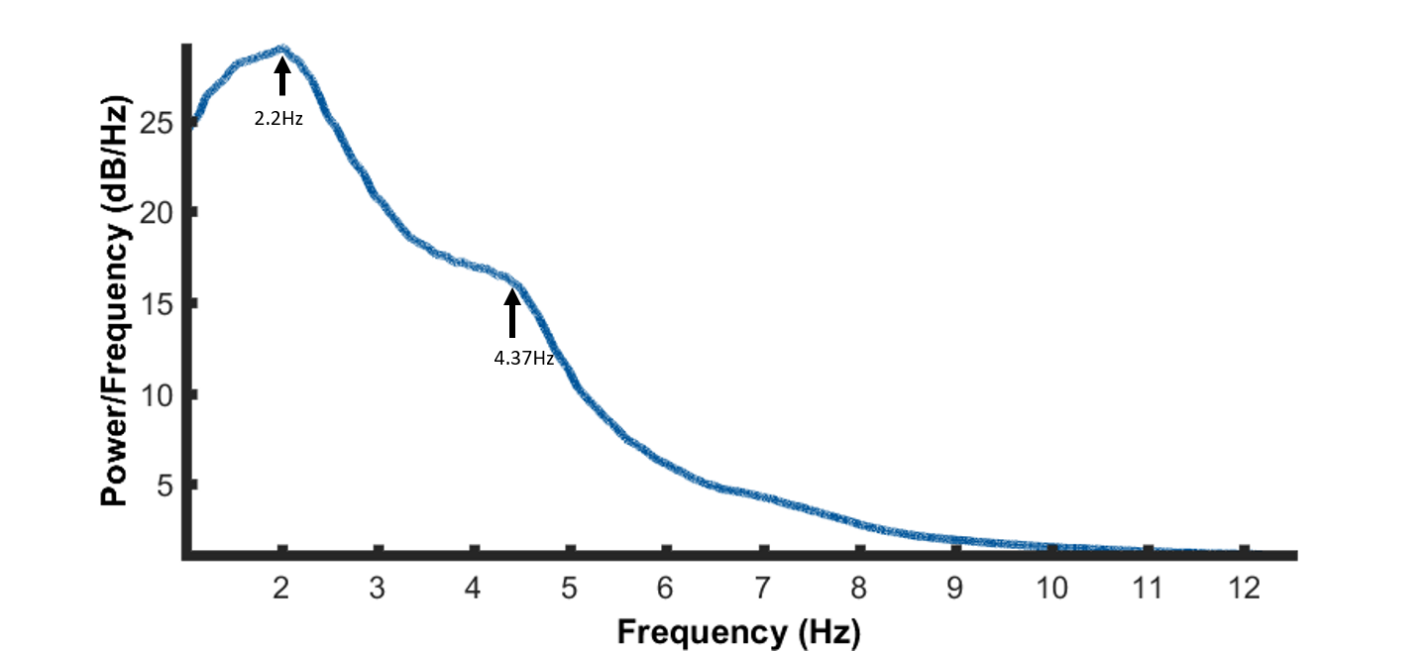

**Figure S3. Grand average spectral decomposition of the EEG signal (1-12 Hz) of the first half of the sample**. A periodogram was used to obtain a power spectral density (PSD) estimate separately for 4-, 7- and 11-months data, before being averaged together to create a grand average. Arrows show the center frequency of peaks of interest. These were defined by visual inspection.

**Modulation Spectrum of nursery rhyme sound files**

All analysis was computed using MATLAB 2016a. The modulation spectrum extraction was based on an approach described by Plomp, et al., (1) and developed by Leong, et al. (2). First, the sound files were down sampled to 14.7 kHz and then band-pass filtered using a series of adjacent FIR filters, into five bands: 100-300 Hz, 300-700 Hz, 700-1750 Hz, 1750-3900 Hz, and 3900- 7250 Hz. Next, the Hilbert envelope was extracted from each of the five sub-band signals. The five envelopes were down sampled to 1050 Hz then filtered through a modulation filter bank. This modulation filter bank comprised 24 channels logarithmically spaced between 0.9-40 Hz. In Figure S4, the RMS difference power was averaged across 18 nursery rhymes and determined for each spectral band. In Figure S5, for one nursery rhyme, the RMS power in each spectral band was divided by the overall RMS power, revealing the relative amount of energy in each band. In Figure S4, the “all band averages” from each of the 18 individual nursery rhymes are depicted together, as denoted by the thin multicolored lines. The grand average of these 18 individual “all band averages” and STD are denoted by the black line and grey shading respectively. Modulation filter bank corner frequencies were taken as [0.93; 1.09; 1.27; 1.49; 1.74; 2.03; 2.38; 2.78; 3.25; 3.80; 4.45; 5.20; 6.08; 7.11; 8.32; 9.72; 11.38; 13.30; 15.56; 18.20; 21.28; 24.89; 29.11; 34.04; 39.81]. Clear peaks in modulation power can be observed at ~2.18 and ~4.4Hz.
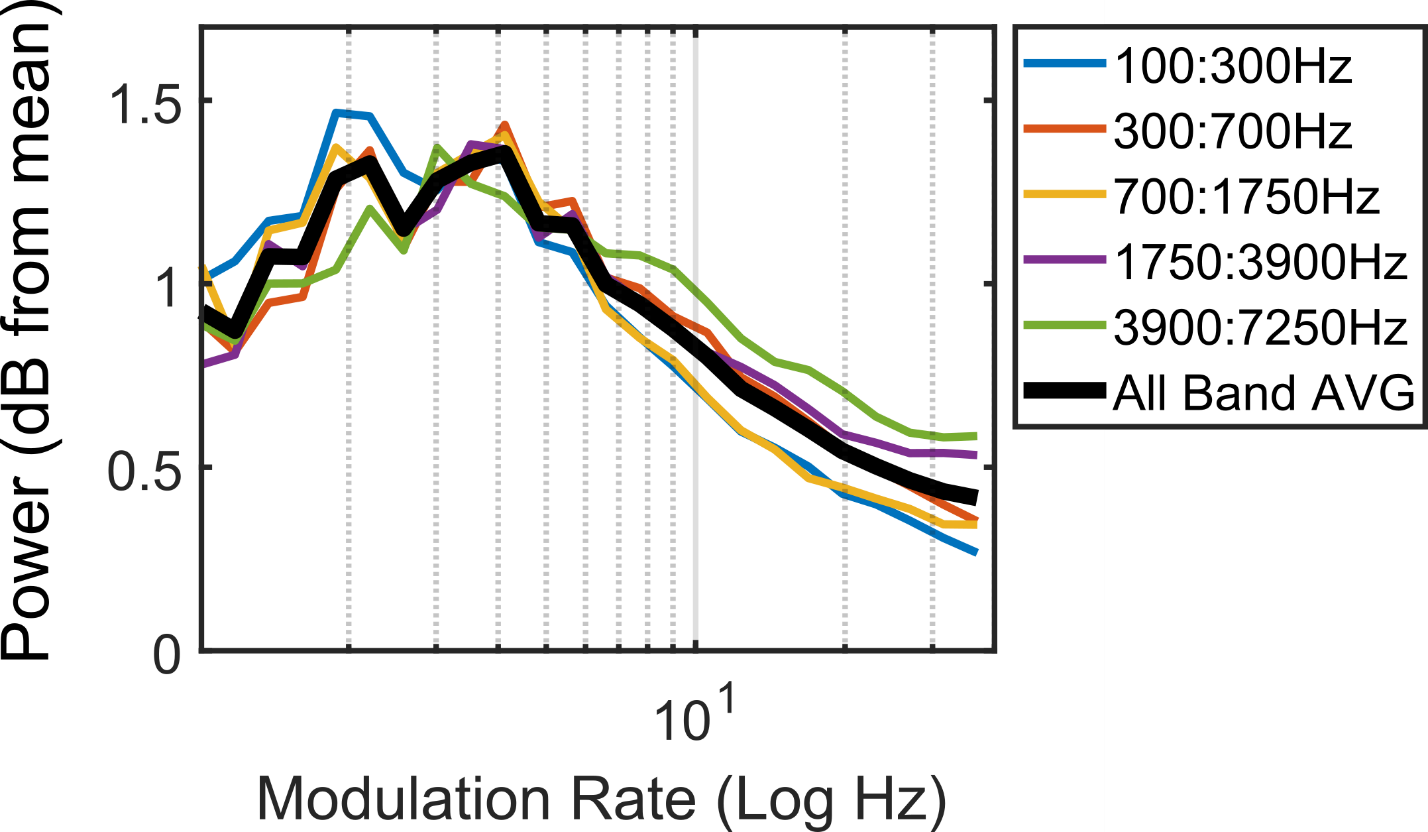

**Figure S4*, Average modulation spectrum of all the Nursery rhyme stimuli.***

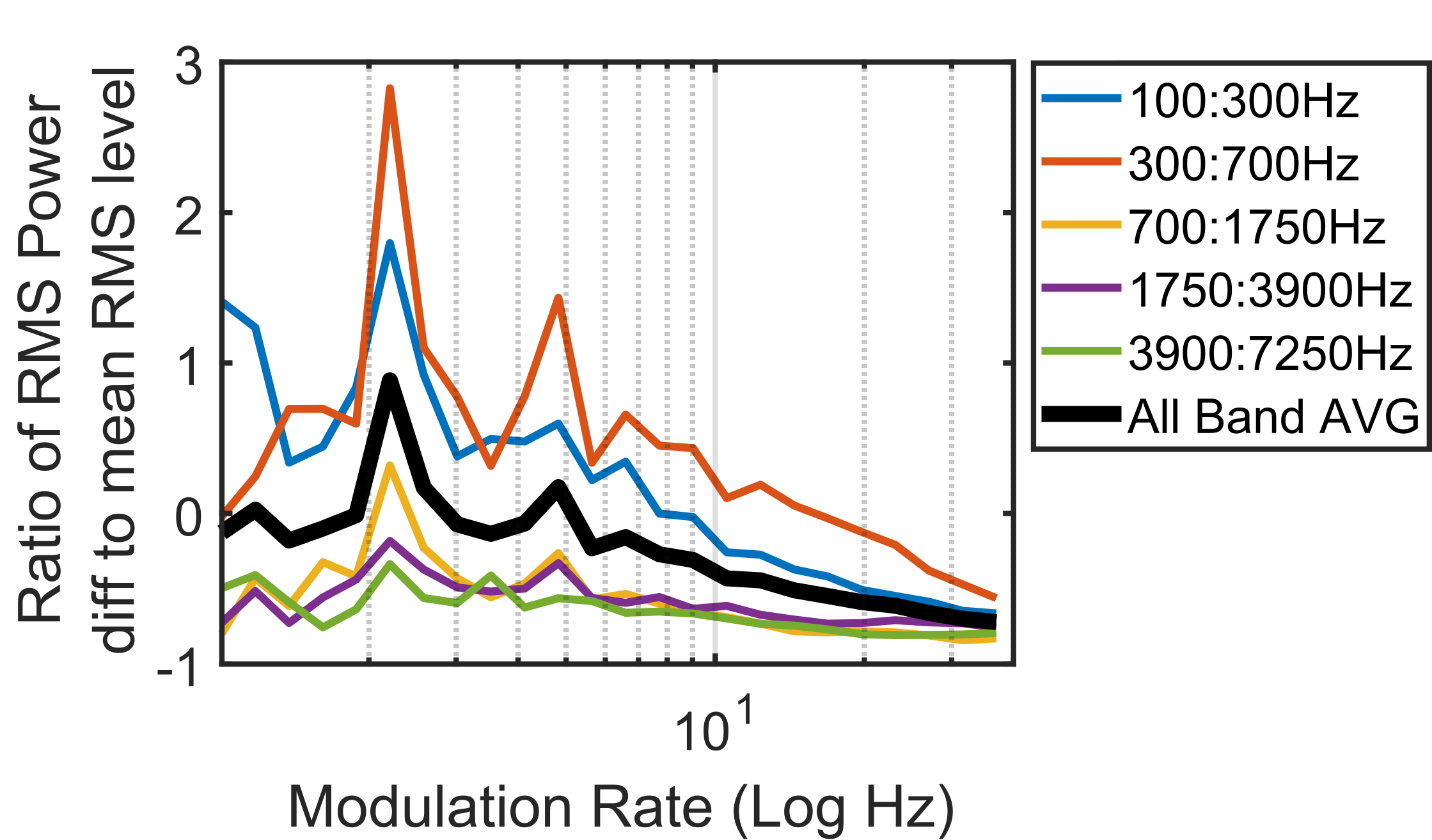

**Figure S5*, Example Modulation spectrum of one the Nursery rhyme stimuli (‘Simple Simon’).*** *RMS power in each spectral band was derived by the overall RMS power.*
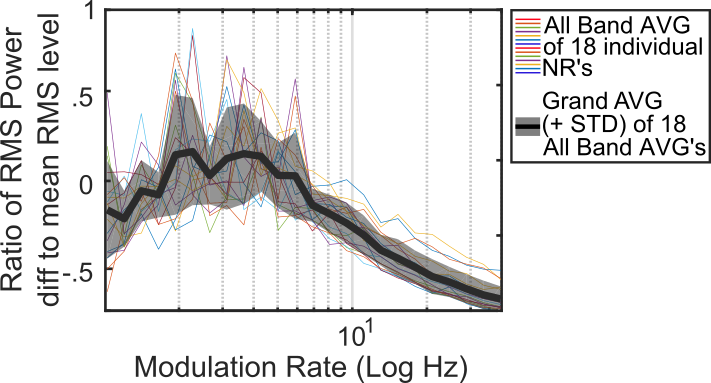

**Figure S6, *Grand average of the 18 individual nursery rhymes “All Band AVG” Modulation spectrums****. The “All Band AVG” is depicted for each individual rhyme by the separate multicolored lines. The grand average of the 18 individual “All Band AVG”, and their standard deviations (STD), are denoted by the black line and grey shading respectively.*

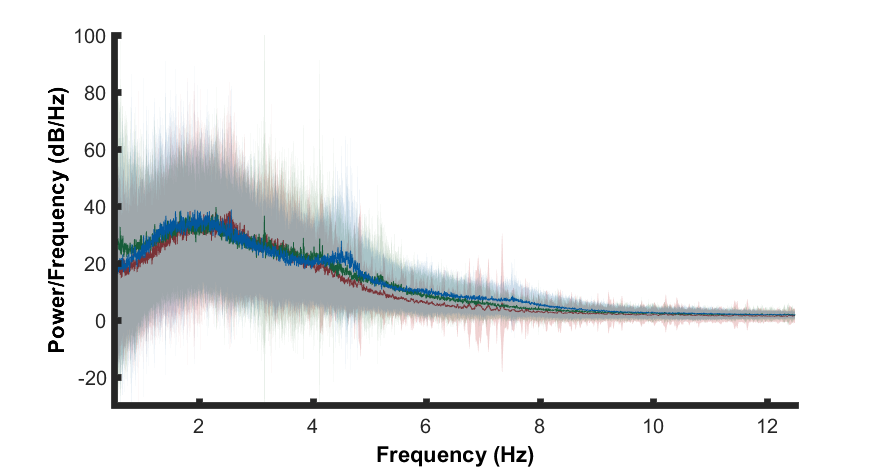

**Figure S7,** **Spectral decomposition of the EEG signal averaged across the full infant cohort (1–12 Hz plotted, from 0.5 to 45 Hz calculated), in response to the silent state period***. A periodogram was used to obtain a power spectral density (PSD) estimate separately for 4- (red), 7- (green) and 11- (blue) months silent state data. First the PSD estimate was averaged across channels per participant. Bold lines indicate the grand average mean values and pale shading plots the standard deviation of the data across the participants. Outlier analysis was also conducted to remove extreme data points that would compromise the LMEM.*

| **a** | *1^st^ half of sample* | | | | | | | *2^nd^ half of sample* | | | | | |
| --- | --- | --- | --- | --- | --- | --- | --- | --- | --- | --- | --- | --- | --- |
| **~1.92 Hz** | *SS* | | | | *NR* | | | *SS* | | | | *NR* | |
|  | *M* | *SE* | | *M* | | *SE* | | *M* | | *SE* | | *M* | *SE* |
| 4 months | 64.57 | 10.49 | | 75.37 | | 9.78 | | 116.25 | | 10.64 | | 97.3 | 10.20 |
| 7 months | 59.08 | 9.96 | | 80.72 | | 9.82 | | 116.33 | | 13.50 | | 124.17 | 13.07 |
| 11 months | 76.73 | 10.10 | | 87.90 | | 9.88 | | 76.67 | | 11.89 | | 94.00 | 11.59 |
|  | *Full sample* | | | | | | | | | | | | |
| **~1.92 Hz** | *SS* | | | | | | | | *NR* | | | | |
|  | *M* | | | | *SE* | | | | *M* | | *SE* | | |
| 4 months | 89.53 | | | | 7.69 | | | | 86.18 | | 7.27 | | |
| 7 months | 82.84 | | | | 8.15 | | | | 100.04 | | 7.95 | | |
| 11 months | 77.27 | | | | 7.86 | | | | 91.33 | | 7.66 | | |
| **b** | *1^st^ half of sample* | | | | | | | *2^nd^ half of sample* | | | | | |
| **~4.05 Hz** | *SS* | | | | *NR* | | | *SS* | | | | *NR* | |
|  | *M* | *SE* | | *M* | | *SE* | | *M* | | *SE* | | *M* | *SE* |
| 4 months | 36.57 | 7.27 | | 34.25 | | 6.34 | | 57.75 | | *6.97* | | 57.63 | 6.28 |
| 7 months | 52.34 | 9.95 | | 65.65 | | 9.75 | | 61.10 | | 8.20 | | 77.49 | 7.47 |
| 11 months | 45.37 | 6.76 | | 61.30 | | 6.47 | | 43.73 | | 7.28 | | 59.96 | 6.79 |
|  | *Full sample* | | | | | | | | | | | | |
| **~4.05 Hz** | *SS* | | | | | | | | *NR* | | | | |
|  | *M* | | | | *SE* | | | | *M* | | *SE* | | |
| 4 months | 45.37 | | | | 4.94 | | | | 45.60 | | 4.34 | | |
| 7 months | 55.61 | | | | 7.07 | | | | 70.66 | | 6.74 | | |
| 11 months | 44.66 | | | | 4.72 | | | | 60.38 | | 4.43 | | |
| **c** | *1^st^ half of sample* | | | | | | | *2^nd^ half of sample* | | | | | |
| **~4.35 Hz** | *SS* | | | | *NR* | | | *SS* | | | | *NR* | |
|  | *M* | *SE* | | *M* | | *SE* | | *M* | | *SE* | | *M* | *SE* |
| 4 months | 28.76 | *7.16* | | 32.13 | | 6.42 | | 47.15 | | *8.35* | | 46.40 | 7.47 |
| 7 months | 43.74 | 8.30 | | 55.19 | | 8.16 | | 59.65 | | 12.74 | | 73.01 | 11.90 |
| 11 months | 49.10 | 7.69 | | 78.26 | | 7.47 | | 56.48 | | 9.46 | | 91.77 | 8.85 |
|  | *Full sample* | | | | | | | | | | | | |
| **~4.35 Hz** | *SS* | | | | | | | | *NR* | | | | |
|  | *M* | | | | *SE* | | | | *M* | | *SE* | | |
| 4 months | 37.94 | | | | 5.47 | | | | 38.89 | | 4.90 | | |
| 7 months | 50.63 | | | | 7.18 | | | | 63.02 | | 6.91 | | |
| 11 months | 53.11 | | | | 6.05 | | | | 84.84 | | 5.78 | | |

***Table S1. Mean and Standard error of the PSD peak amplitudes, for both recording Silent state and Nursery rhyme recording conditions.*** *Table (a) provides the average of the peak amplitudes (across all subjects) within a 0.25 Hz window centred on 1.92 Hz. The data are separated into the first and second half of the sample, and also given for the full sample. Table (b) provides the average of the peak amplitudes (across all subjects) within a 0.25 Hz window cantered on 4.05 Hz. The data are separated into the first and second half of the sample and also given for the full sample. Table (c) provides the average of the peak amplitudes (across all subjects) within a 0.25 Hz window cantered on 4.35 Hz. The data are separated into the first and second half of the sample, and also given for the full sample.*

### *PSD estimates of fixed effects*

| **PSD estimates of fixed effects (full sample)** |
| --- |

| a) **1.92 Hz** |  |  |  |  |  |
| --- | --- | --- | --- | --- | --- |
| *full sample* | *β est.* | *SE_b_* | *df* | *p* | *95% CI* |
| Recording type (stim. rel. to silent) | 14.066 | 5.479 | 189.23 | 0.011 | [3.259 24.873] |
| Age 4mo (rel. 11mo) | 12.261 | 9.136 | 438.83 | 0.180 | [-5.696 30.218] |
| Age 7mo (rel. 11mo) | 5.572 | 9.459 | 273.73 | 0.556 | [-13.050 24.194] |
| Recording type * 4mo (rel. 11mo) | -17.411 | 7.857 | 191.76 | 0.028 | [ -32.909 -1.913] |
| Recording type * 7mo (rel. 11mo) | 3.136 | 7.758 | 185.90 | 0.686 | [-12.169 18.442] |

| b) **4.05 Hz** |  |  |  |  |  |
| --- | --- | --- | --- | --- | --- |
| *Full sample* | *β est.* | *SE_b_* | *df* | *p* | *95% CI* |
| Recording type (stim. rel. to silent) | 15.722 | 6.477 | 416.55 | .016 | [2.990 28.454] |
| Age 4mo (rel. 11mo) | 0.718 | 6.836 | 416.55 | .916 | [-12.720 14.156] |
| Age 7mo (rel. 11mo) | 10.950 | 8.504 | 283.07 | .199 | [-5.788 27.688] |
| Recording type * 4mo (rel. 11mo) | -15.495 | 9.230 | 416.55 | .094 | [-33.639 2.649] |
| Recording type * 7mo (rel. 11mo) | -0.671 | 9.375 | 428.19 | .943 | [-19.100 17.756] |
| c) **4.35 Hz** |  |  |  |  |  |
| *Full sample* | *β est.* | *SE_b_* | *df* | *p* | *95% CI* |
| Recording type (stim. rel. to silent) | 31.730 | 5.876 | 238.71 | 1.6x10^-7^ | [20.154 43.305] |
| Age 4mo (rel. 11mo) | -15.167 | 7.434 | 430.30 | .042 | [-29.778 0.556] |
| Age 7mo (rel. 11mo) | -2.479 | 8.738 | 269.70 | .777 | [-19.682 14.725] |
| Recording type * 4mo (rel. 11mo) | -30.783 | 8.355 | 241.59 | 2.8x10^-4^ | [-47.242 -14.325] |
| Recording type * 7mo (rel. 11mo) | -19.339 | 8.372 | 232.22 | .022 | [-35.833 -2.844] |

**Table S2. Parameter estimates from the full sample PSD LMEM.** Detailing estimates of fixed effects on periodogram PSD peaks at a) 1.92 Hz and at b, 4.05 Hz and c) 4.35 Hz, taken from the LMEM described in the text. The base cases for the LMEM were set as 11-months for age and silent state for recording type.

### *Cortical tracking estimates of fixed effects*

| **Cortical tracking estimates of fixed effects (full sample)** |  |  |  |  |  |
| --- | --- | --- | --- | --- | --- |
| *Full sample* | *β _estimate_* | *SE_b_* | *df* | *p* | *95% CI* |
| Data type (real rel. to rand for α base case) | -5.1x10^-5^ | 0.001 | 1625 | 0.960 | [-0.002 0.001] |
| Delta (rel. to α) | 0.005 | 0.001 | 1623.64 | 2.0x10^-6^ | [0.008 0.0021] |
| Theta (rel. to α) | 0.004 | 0.001 | 1629.35 | 9.5x10^-4^ | [0.001 0.006] |
| Age (4mo rel. to 11mo) | -0.001 | 0.001 | 1869.63 | 0.302 | [-0.003 0.0001] |
| Age (7mo rel. to 11mo) | 4.8x10^-6^ | 0.001 | 1741.17 | 0.997 | [-0.002 0.002] |
| Data type * delta (rel. to α) | 0.015 | 0.001 | 1624.10 | 9.0x10^-40^ | [0.013 0.017] |
| Data type * theta (rel. to α) | 0.004 | 0.001 | 1625.85 | 6.8x10^-5^ | [0.002 0.007] |
| Data type * 4mo (rel. to 11mo) | 0.002 | 0.001 | 1625.19 | 0.101 | [0.0004 0.004] |
| Data type * 7mo (rel. to 11mo) | -5.7x10^-4^ | 0.001 | 1625.92 | 0.609 | [-0.003 0.002] |
| 4mo * delta (rel. to α) | 0.006 | 0.001 | 1625.66 | 4.5x10^-5^ | [0.003 0.008] |
| 4mo * theta (rel. to α) | 0.003 | 0.001 | 1626.15 | 0.045 | [5.7x10^-5^ 0.005] |
| 7mo * delta (rel. to α) | -1.4x10^-4^ | 0.001 | 1627.80 | 0.917 | [-0.003 0.003] |
| 7mo * theta (rel. to α) | 2.0x10^-4^ | 0.001 | 1628.76 | 0.885 | [-0.002 0.003] |

**Table S3. Parameter estimates from the full sample cortical tracking LMEM**. Individual subject Pearson correlations (r values) derived from 3 frequency bands (delta 0.5–4 Hz, theta 4–8 Hz and alpha 8–12 Hz), 3 ages (4-, 7- or 11-months) and 2 data types (random or real). Random permutation r values, alpha and 11-months data were set as the respective base cases for data type, frequency band and age in the model.

### *Phase amplitude coupling (PAC) estimates of fixed effects*

| **PAC estimates of fixed effects** |  |  |  |  |  |
| --- | --- | --- | --- | --- | --- |
| *Full sample* | *β _est._* | *SE_b_* | *df* | *p* | *95% CI* |
| Data type (random re. to real nMI) | -1.296 | 0.033 | 2269.31 | ***<.001*** | [-1.361 -1.231] |
| Low-frequency phase (rel. to δ) | .370 | .039 | 2269.31 | ***<.001*** | [.293 .447] |
| High-frequency amplitude (rel. to γ) | .036 | .039 | 2269.31 | .360 | [-.041 .113] |
| Age (11mo rel. to 4mo) | .020 | .041 | 2523.35 | .624 | [-.061 .102] |
| Age (7mo rel. to 4mo) | -.048 | .042 | 2524.69 | .248 | [-.131 .034] |
| Data type * LFP | -.320 | .030 | 2269.31 | ***<.001*** | [-.379 -.262] |
| Data type * HFA | -.040 | .030 | 2269.31 | .182 | [-.098 .019] |
| Data type * 11mo (rel. to 4mo) | .061 | .036 | 2269.31 | .091 | [-.010 .132] |
| Data type * 7mo (rel. to 4mo) | .073 | .037 | 2269.31 | .047 | [.001 .145] |
| LFP * 11mo (rel. to 4mo) | -.095 | .051 | 2269.31 | .065 | [-.195 .006] |
| LFP * 7mo (rel. to 4mo) | -.084 | .052 | 2269.31 | .104 | [-.186 .017] |
| HFA * 11mo (rel. to 4mo) | -.047 | .051 | 2269.31 | .365 | [-.147 .054] |
| HFA * 7mo (rel. to 4mo) | .008 | .052 | 2269.31 | .883 | [-.094 .109] |
| HFA*LFP | -.011 | .051 | 2269.31 | .833 | [-.111 .090] |
| HFA*LFP*11mo (rel. to 4mo) | .039 | .073 | 2269.31 | 0.588 | [-.103 .182] |
| HFA*LFP*7mo (rel. to 4mo) | .069 | .073 | 2269.31 | 0.348 | [-.075 .212] |

**Table S4. Parameter estimates from the full sample PAC LMEM**. The normalised modulation index (nMI) measure of PAC was calculated for two data types (real and random nMI values), between multiple low frequency phases (LFP) and High frequency amplitude (HFA) calculation pairs (LFP steps 2:8 Hz and HFA steps from 15:45 Hz). The maximum nMI value was taken from four pre-defined PAC band groupings (delta/beta, delta/gamma, theta/beta and theta/gamma; beta 2–4 Hz, theta 4–8 Hz, beta 15–30 Hz and gamma 30–45 Hz) and was submitted to the model. The base cases for the LMEM were set as real nMI data for data type, 4-months for age, delta (δ) for LFP and gamma (**γ**) for HFA.

### *~1.92Hz PSD power as a predictor of language acquisitions*

A multivariate linear model was conducted to see if ~1.92Hz PSD power at 4- or 7-months predicted either infant-led or parent-estimated language outcomes. Posthoc univariate linear models are also reported in Table S5 & S6 to demonstrate the individual contribution of each language measure to global parent-estimated or infant-led language performance.

| **~1.92 Hz PSD power at 4 months**  **a, Infant-led language measures** |  |  |  |  |
| --- | --- | --- | --- | --- |
| **Multivariate linear model** | *Df _num_* | *Df _denom_* | *F* | *p* |
| Infant-led language measures | 5 | 68 | 0.41 | 0.84 |
| **Univariate linear models (post hoc)** | *Df* | *β _estimate_* | *SE* | *P* |
| NWR Consonants, 24 months | 81 | 1.9 x10^-4^ | 4.6 x10^-4^ | 0.69 |
| NWR Syllables, 24 months | 81 | -9.0 x10^-4^ | 4.5 x10^-4^ | 0.84 |
| NWR Stress, 24 months | 81 | 4.3 x10^-4^ | 4.3 x10^-4^ | 0.33 |
| CCT, 18 months | 81 | 2.3 x10^-4^ | 4.3 x10^-4^ | 0.50 |
| Pointing, 12 months | 89 | 4.3 x10^-4^ | 7.5 x10^-4^ | 0.56 |
| **b, Parent-estimated language measures** |  |  |  |  |
| **Multivariate linear model** | *Df _num_* | *Df _demom_* | *F* | *p* |
| Parent-estimated language measures | 2 | 76 | 1.58 | 0.21 |
| **Univariate linear models (post hoc)** | *Df* | *β _estimate_* | *SE* | *p* |
| CDI comprehension, 24 months | 79 | 0.30 | 0.22 | 0.18 |
| CDI production, 24 months | 79 | 0.24 | 0.24 | 0.08 |

**Table S5. Multivariate and univariate linear models for 1.92 Hz PSD power at 4 months.** Investigating whether ~1.92 Hz PSD power at 4 months predicted either infant-led or parent-estimated language outcomes. The table details the multivariate linear models (reporting the global effects) and the univariate linear models (ran after the main multivariate model) describing whether ~1.92 Hz PSD power at 4 months predicted a) infant-led language measures or b) the parent-estimated language measures. Bonferroni correction for multiple comparisons in the **univariate models** led to modified significant alpha levels of, p = <0.01 for infant-led and p = <0.025 for parent-estimated.

| **~1.92 Hz PSD power at 7 months**  **a, Infant-led language measures** |  |  |  |  |
| --- | --- | --- | --- | --- |
| **Multivariate linear model** | *Df _num_* | *Df _denom_* | *F* | *p* |
| Infant-led language measures | 5 | 66 | 1.02 | 0.41 |
| **Univariate linear models (post hoc)** | *Df* | *β _estimate_* | *SE* | *p* |
| NWR Consonants, 24 months | 1,80 | -4.2 x10^-4^ | 4.6 x10^-4^ | 0.37 |
| NWR Syllables, 24 months | 1,80 | -3.6 x10^-4^ | 4.6 x10^-4^ | 0.43 |
| NWR Stress, 24 months | 1,80 | 9.4 x10^-4^ | 4.0 x10^-4^ | 0.98 |
| CCT, 18 months | 1,76 | 2.0 x10^-4^ | 3.4 x10^-4^ | 0.57 |
| Pointing, 12 months | 1,85 | 4.8 x10^-4^ | 7.4 x10^-4^ | 0.52 |
| **b, Parent-estimated language measures** |  |  |  |  |
| **Multivariate linear model** | *Df _num_* | *Df _denom_* | *F* | *p* |
| Parent-estimated language measures | 2 | 73 | 0.16 | 0.85 |
| **Univariate linear models (post hoc)** | *Df* | *β _estimate_* | *SE* | *p* |
| CDI comprehension, 24 months | 1,74 | 0.10 | 0.23 | 0.66 |
| CDI production, 24 months | 1,74 | 0.04 | 0.24 | 0.88 |

**Table S6. Multivariate and univariate linear models for ~1.92 Hz PSD power at 7 months.** Investigating whether ~1.92 Hz PSD power at 7 months predicted either infant-led or parent-estimated language outcomes. The table details the multivariate linear models (reporting the global effects) and the univariate linear models (ran after the main multivariate model) describing whether ~1.92 Hz PSD power at 7 months predicted a) infant-led language measures or b) the parent estimated language measures. Bonferroni correction for multiple comparisons in the **univariate models** led to modified significant alpha levels of, p = <0.01 for infant-led and p = <0.025 for parent-estimated.

### *~4.05Hz PSD power as a predictor of language acquisition*

A multivariate linear model was conducted to see if ~4.05 Hz PSD power at 4-, 7 or 11-months predicted either infant-led or parent-estimated language outcomes. Posthoc univariate linear models are also reported in Tables S7-S9 to demonstrate the individual contribution of each language measure to global parent-estimated or infant-led language performance.

| **~4.05 Hz PSD power at 4 months**  **a, Infant-led language measures** |  |  |  |  |
| --- | --- | --- | --- | --- |
| **Multivariate linear model** | *Df _num_* | *Df _denom_* | *F* | *p* |
| Infant-led language measures | 5 | 68 | 0.31 | 0.91 |
| **Univariate linear models (post hoc)** | *Df* | *β _estimate_* | *SE* | *P* |
| NWR Consonants, 24 months | 81 | 4.2 x10^-4^ | 7.5 x10^-4^ | 0.58 |
| NWR Syllables, 24 months | 81 | -1.1 x10^-4^ | 7.3 x10^-4^ | 0.88 |
| NWR Stress, 24 months | 81 | 5.8 x10^-4^ | 7.0 x10^-4^ | 0.41 |
| CCT, 18 months | 81 | 1.5 x10^-4^ | 6.9 x10^-4^ | 0.99 |
| Pointing, 12 months | 89 | -0.001 | 0.001 | 0.37 |
| **b, Parent-estimated language measures** |  |  |  |  |
| **Multivariate linear model** | *Df _num_* | *Df _denom_* | *F* | *p* |
| Parent-estimated language measures | 2 | 76 | 2.42 | 0.96 |
| **Univariate linear models (post hoc)** | *Df* | *β _estimate_* | *SE* | *p* |
| CDI comprehension, 24 months | 79 | 0.76 | 0.35 | **0.03** |
| CDI production, 24 months | 79 | 0.59 | 0.38 | 0.13 |

**Table S7. Multivariate and univariate linear models for ~4.05 Hz PSD power at 4 months.** Investigating whether ~4.05 Hz PSD power at 4 months predicted either infant-led or parent-estimated language outcomes. The table details the multivariate linear models (reporting the global effects) and the univariate linear models (ran after the main multivariate model) describing whether ~4.05 Hz PSD power at 4 months predicted a) infant-led language measures or b) the parent estimated language measures. Bonferroni correction for multiple comparisons in the **univariate models** led to modified significant alpha levels of, p = <0.01 for infant-led and p = <0.025 for parent-estimated.

| **~4.05 Hz PSD power at 7 months**  **a, Infant-led language measures** |  |  |  |  |
| --- | --- | --- | --- | --- |
| **Multivariate linear model** | *Df _num_* | *Df _denom_* | *F* | *p* |
| Infant-led language measures | 5 | 66 | 0.25 | 0.94 |
| **Univariate linear models (post hoc)** | *Df* | *β _estimate_* | *SE* | *p* |
| NWR Consonants, 24 months | 80 | -3.8 x10^-4^ | 5.2 x10^-4^ | 0.46 |
| NWR Syllables, 24 months | 80 | -6.0 x10^-4^ | 5.1 x10^-4^ | 0.24 |
| NWR Stress, 24 months | 80 | -3.9 x10^-4^ | 4.5 x10^-4^ | 0.38 |
| CCT, 18 months | 76 | 1.8 x10^-4^ | 3.9 x10^-4^ | 0.64 |
| Pointing, 12 months | 85 | 5.9 x10^-4^ | 8.4 x10^-4^ | 0.94 |
| **b, Parent-estimated language measures** |  |  |  |  |
| **Multivariate linear model** | *Df _num_* | *Df _denom_* | *F* | *p* |
| Parent-estimated language measures | 2 | 73 | 1.88 | 0.16 |
| **Univariate linear models (post hoc)** | *Df* | *β _estimate_* | *SE* | *p* |
| CDI comprehension, 24 months | 74 | 0.25 | 0.96 | 0.34 |
| CDI production, 24 months | 74 | 0.27 | -0.15 | 0.88 |

**Table S8. Multivariate and univariate linear models for ~4.05 Hz PSD power at 7 months.** Investigating whether ~4.05 Hz PSD power at 7 months predicted either infant-led or parent-estimated language outcomes. The table details the multivariate linear models (reporting the global effects) and the univariate linear models (ran after the main multivariate model) describing whether ~4.05 Hz PSD power at 7 months predicted a) infant-led language measures or b) the parent estimated language measures. Bonferroni correction for multiple comparisons in the **univariate models** led to modified significant alpha levels of, p = <0.01 for infant-led and p = <0.025 for parent-estimated.

| **~4.05 Hz PSD power at 11 months**  **a, Infant-led language measures** |  |  |  |  |
| --- | --- | --- | --- | --- |
| **Multivariate linear model** | *Df _num_* | *Df _denom_* | *F* | *p* |
| Infant-led language measures | 5 | 65 | 1.43 | 0.22 |
| **Univariate linear models (post hoc)** | *Df* | *β _estimate_* | *SE* | *p* |
| NWR Consonants, 24 months | 79 | 2.4 x10^-4^ | 9.0 x10^-4^ | 0.79 |
| NWR Syllables, 24 months | 79 | -6.9 x10^-4^ | 9.1 x10^-4^ | 0.45 |
| NWR Stress, 24 months | 79 | -1.9 x10^-4^ | 8.3 x10^-4^ | 0.82 |
| CCT, 18 months | 79 | 0.001 | 6.7 x10^-4^ | 0.12 |
| Pointing, 12 months | 90 | 0.002 | 0.001 | 0.29 |
| **b, Parent-estimated language measures** |  |  |  |  |
| **Multivariate linear model** | *Df _num_* | *Df _denom_* | *F* | *p* |
| Parent-estimated language measures | 2 | 75 | 0.84 | 0.44 |
| **Univariate linear models (post hoc)** | *Df* | *β _estimate_* | *SE* | *p* |
| CDI comprehension, 24 months | 78 | 0.332 | 0.464 | 0.476 |
| CDI production, 24 months | 78 | 0.030 | 0.484 | 0.951 |

**Table S9. Multivariate and univariate linear models for ~4.05 Hz PSD power at 11 months.** Investigating whether ~4.05 Hz PSD power at 11 months predicted either infant-led or parent-estimated language outcomes. The table details the multivariate linear models (reporting the global effects) and the univariate linear models (ran after the main multivariate model) describing whether ~4.05 Hz PSD power at 11 months predicted a) infant-led language measures or b) the parent estimated language measures. Bonferroni correction for multiple comparisons in the **univariate models** led to modified significant alpha levels of, p = <0.01 for infant-led and p = <0.025 for parent-estimated.

### *~4.35Hz PSD power as a predictor of language acquisitions*

A multivariate linear model was conducted to see if ~4.35 Hz PSD power at 4-, or 7-months predicted either infant-led or parent-estimated language outcomes. Posthoc univariate linear models are also reported in Tables S10 & S11 to demonstrate the individual contribution of each language measure to global parent-estimated or infant-led language performance.

| **~4.35 Hz PSD power at 4 months**  **a, Infant-led language measures** |  |  |  |  |
| --- | --- | --- | --- | --- |
| **Multivariate linear model** | *Df _num_* | *Df _denom_* | *F* | *p* |
| Infant-led language measures | 5 | 68 | 0.18 | 0.97 |
| **Univariate linear models (post hoc)** | *Df* | *β _estimate_* | *SE* | *p* |
| NWR Consonants, 24 months | 81 | 6.6 x10^-4^ | 0.001 | 0.56 |
| NWR Syllables, 24 months | 81 | -4.6 x10^-4^ | 1.1 x10^-4^ | 0.97 |
| NWR Stress, 24 months | 81 | 0.001 | 0.001 | 0.28 |
| CCT, 18 months | 81 | 2.3 x10^-4^ | 9.8 x10^-4^ | 0.82 |
| Pointing, 12 months | 89 | -0.001 | 0.002 | 0.46 |
| **b, Parent-estimated language measures** |  |  |  |  |
| **Multivariate linear model** | *Df _num_* | *Df _denom_* | *F* | *p* |
| Parent-estimated language measures | 2 | 76 | 1.87 | 0.16 |
| **Univariate linear models (post hoc)** | *Df* | *β _estimate_* | *SE* | *p* |
| CDI comprehension, 24 months | 79 | 1.04 | 0.53 | 0.06 |
| CDI production, 24 months | 79 | 0.88 | 0.58 | 0.13 |

**Table S10. Multivariate and univariate linear models for ~4.35 Hz PSD power at 4 months.** Investigating whether ~4.35 Hz PSD power at 4 months predicted either infant-led or parent-estimated language outcomes. The table details the multivariate linear models (reporting the global effects) and the univariate linear models (ran after the main multivariate model) describing whether ~4.35 Hz PSD power at 4 months predicted a) infant-led language measures or b) the parent estimated language measures. Bonferroni correction for multiple comparisons in the **univariate models** led to modified significant alpha levels of, p = <0.01 for infant-led and p = <0.025 for parent-estimated.

| **~4.35 Hz PSD power at 7 months**  **a, Infant-led language measures** |  |  |  |  |
| --- | --- | --- | --- | --- |
| **Multivariate linear model** | *Df _num_* | *Df _denom_* | *F* | *p* |
| Infant-led language measures | 5 | 66 | 1.36 | 0.25 |
| **Univariate linear models (post hoc)** | *Df* | *β _estimate_* | *SE* | *p* |
| NWR Consonants, 24 months | 1,80 | 8.4 x10^-4^ | 6.2 x10^-4^ | 0.18 |
| NWR Syllables, 24 months | 1,80 | -0.001 | 6.2 x10^-4^ | 0.11 |
| NWR Stress, 24 months | 1,80 | -4.0 x10^-4^ | 5.4 x10^-4^ | 0.46 |
| CCT, 18 months | 1,76 | -3.1 x10^-4^ | 5.4 x10^-4^ | 0.56 |
| Pointing, 12 months | 1,85 | -4.8 x10^-4^ | 0.001 | 0.63 |
| **b, Parent-estimated language measures** |  |  |  |  |
| **Multivariate linear model** | *Df _num_* | *Df _denom_* | *F* | *p* |
| Parent-estimated language measures | 2 | 73 | 0.46 | 0.63 |
| **Univariate linear models (post hoc)** | *Df* | *β _estimate_* | *SE* | *p* |
| CDI comprehension, 24 months | 1,76 | -0.22 | 0.31 | 0.48 |
| CDI production, 24 months | 1,76 | -0.32 | 0.33 | 0.34 |

**Table S11. Multivariate and univariate linear models for ~4.35 Hz PSD power at 7 months.** Investigating whether ~4.35 Hz PSD power at 7 months predicted either infant-led or parent-estimated language outcomes. The table details the multivariate linear models (reporting the global effects) and the univariate linear models (ran after the main multivariate model) describing whether ~4.35 Hz PSD power at 7 months predicted a) infant-led language measures or b) the parent estimated language measures. Bonferroni correction for multiple comparisons in the **univariate models** led to modified significant alpha levels of, p = <0.01 for infant-led and p = <0.025 for parent-estimated*.*

### Delta cortical tracking as a predictor of language acquisition

A multivariate linear model was conducted to see if delta cortical tracking at 4- or 7-months predicted either infant-led or parent-estimated language outcomes. Posthoc univariate linear models are also reported in Tables S12 & S13 to demonstrate the individual contribution of each language measure to global parent-estimated or infant-led language performance.

| **Delta cortical tracking at 4mo**  **a, Infant-led language measures** |  |  |  |  |
| --- | --- | --- | --- | --- |
| **Multivariate linear model** | *Df _num_* | *Df _denom_* | *F* | *p* |
| Infant-led language measures | 5 | 75 | 0.96 | 0.45 |
| **Univariate linear models (post hoc)** | *Df* | *β _estimate_* | *SE* | *p* |
| NWR Consonants, 24 months | 1,93 | -0.41 | 1.82 | 0.82 |
| NWR Syllables, 24 months | 1,93 | -0.29 | 1.82 | 0.87 |
| NWR Stress, 24 months | 1,93 | 1.57 | 1.68 | 0.35 |
| CCT, 18 months | 1,92 | -0.20 | 1.13 | 0.88 |
| Pointing, 12 months | 1,103 | 4.49 | 2.59 | 0.09 |
| **b, Parent-estimated language measures** |  |  |  |  |
| **Multivariate linear model** | *Df _num_* | *Df _denom_* | *F* | *p* |
| Parent-estimated language measures | 2 | 84 | 0.99 | 0.38 |
| **Univariate linear models (post hoc)** | *Df* | *β _estimate_* | *SE* | *p* |
| CDI comprehension, 24 months | 1,87 | -1169.27 | 842.55 | 0.17 |
| CDI production, 24 months | 1,87 | -903.90 | 921.79 | 0.33 |

**Table S12. *Multivariate and univariate linear models for delta cortical tracking at 4 months.*** *Investigating whether delta cortical tracking at 4 months predicted either infant-led or parent-estimated language outcomes. The table details the multivariate linear models (reporting the global effects) and the univariate linear models (ran after the main multivariate model) describing whether delta cortical tracking at 4 months predicted a) infant-led language measures or b) the parent estimated language measures. Bonferroni correction for multiple comparisons in the* ***univariate models*** *led to modified significant alpha levels of, p = <0.01 for infant-led and p = <0.025 for parent-estimated (denoted by* ***bold italic text****).*

| **Delta cortical tracking at 7mo**  **a, Infant-led language measures** |  |  |  |  |
| --- | --- | --- | --- | --- |
| **Multivariate linear model** | *Df _num_* | *Df _denom_* | *F* | *p* |
| Infant-led language measures | 5 | 76 | 0.89 | 0.49 |
| **Univariate linear models (post hoc)** | *Df* | *β _estimate_* | *SE* | *p* |
| NWR Consonants, 24 months | 1,95 | 1.3 | 1.68 | 0.44 |
| NWR Syllables, 24 months | 1,95 | 1.61 | 1.66 | 0.34 |
| NWR Stress, 24 months | 1,95 | 0.55 | 1.57 | 0.73 |
| CCT, 18 months | 1,90 | 2.21 | 1.31 | 0.09 |
| Pointing, 12 months | 1,102 | -1.94 | 2.72 | 0.48 |
| **b, Parent-estimated language measures** |  |  |  |  |
| **Multivariate linear model** | *Df _num_* | *Df _denom_* | *F* | *p* |
| Parent-estimated language measures | 2 | 85 | 2.63 | **0.08** |
| **Univariate linear models (post hoc)** | *Df* | *β _estimate_* | *SE* | *p* |
| CDI comprehension, 24 months | 1,88 | -1562.89 | 886.37 | 0.08 |
| CDI production, 24 months | 1,88 | -593.12 | 967.22 | 0.54 |

**Table S13. *Multivariate and univariate linear models for delta cortical tracking at 7 months.*** *Investigating whether delta cortical tracking at 7 months predicted either infant-led or parent-estimated language outcomes. The table details the multivariate linear models (reporting the global effects) and the univariate linear models (ran after the main multivariate model) describing whether delta cortical tracking at 7 months predicted a) infant-led language measures or b) the parent estimated language measures. Bonferroni correction for multiple comparisons in the* ***univariate models*** *led to modified significant alpha levels of, p = <0.01 for infant-led and p = <0.025 for parent-estimated (denoted by* ***bold italic text****).*

### Theta cortical tracking as a predictor of language acquisition

A multivariate linear model was conducted to see if theta cortical tracking at 4- or 7-months predicted either infant-led or parent-estimated language outcomes. Posthoc univariate linear models are also reported in Tables S14 & S15 to demonstrate the individual contribution of each language measure to global parent-estimated or infant-led language performance.

| **Theta cortical tracking 4mo**  **a, Infant-led language measures** |  |  |  |  |
| --- | --- | --- | --- | --- |
| **Multivariate linear model** | *Df _num_* | *Df _denom_* | *F* | *p* |
| Infant-led language measures | 5 | 75 | 0.83 | 0.53 |
| **Univariate linear models (post hoc)** | *Df* | *β _estimate_* | *SE* | *p* |
| NWR Consonants, 24 months | 1,93 | 0.39 | 3.21 | 0.90 |
| NWR Syllables, 24 months | 1,93 | 1.48 | 3.19 | 0.65 |
| NWR Stress, 24 months | 1,93 | 0.10 | 2.97 | 0.97 |
| CCT, 18 months | 1,92 | 3.69 | 2.25 | 0.11 |
| Pointing, 12 months | 1,103 | 4.41 | 4.86 | 0.37 |
| **b, Parent-estimated language measures** |  |  |  |  |
| **Multivariate linear model** | *Df _num_* | *Df _denom_* | *F* | *p* |
| Parent-estimated language measures | 2 | 84 | 0.15 | 0.86 |
| **Univariate linear models (post hoc)** | *Df* | *β _estimate_* | *SE* | *p* |
| CDI comprehension, 24 months | 1,87 | 853.87 | 1702.98 | 0.62 |
| CDI production, 24 months | 1,87 | 1009.37 | 1862.17 | 0.59 |

**Table S14. *Multivariate and univariate linear models for theta cortical tracking at 4 months.*** *Investigating whether Theta cortical tracking at 4 months predicted either infant-led or parent-estimated language outcomes. The table details the multivariate linear models (reporting the global effects) and the univariate linear models (ran after the main multivariate model) describing whether theta cortical tracking at 4 months predicted a) infant-led language measures or b) the parent estimated language measures. Bonferroni correction for multiple comparisons in the* ***univariate models*** *led to modified significant alpha levels of, p = <0.01 for infant-led and p = <0.025 for parent-estimated (denoted by* ***bold italic text****).*

| **Theta cortical tracking at 7mo**  **a, Infant-led language measures** |  |  |  |  |
| --- | --- | --- | --- | --- |
| **Multivariate linear model** | *Df _num_* | *Df _denom_* | *F* | *p* |
| Infant-led language measures | 5 | 76 | 1.31 | 0.27 |
| **Univariate linear models (post hoc)** | *Df* | *β _estimate_* | *SE* | *p* |
| NWR Consonants, 24 months | 95 | 1.28 | 3.08 | 0.68 |
| NWR Syllables, 24 months | 95 | 1.41 | 3.04 | 0.65 |
| NWR Stress, 24 months | 95 | 2.00 | 2.86 | 0.49 |
| CCT, 18 months | 90 | -5.20 | 2.19 | 0.02 |
| Pointing, 12 months | 102 | -7.74 | 4.80 | 0.11 |
| **b, Parent-estimated language measures** |  |  |  |  |
| **Multivariate linear model** | *Df _num_* | *Df _denom_* | *F* | *p* |
| Parent-estimated language measures | 2 | 85 | 0.17 | 0.84 |
| **Univariate linear models (post hoc)** | *Df* | *β _estimate_* | *SE* | *p* |
| CDI comprehension, 24 months | 88 | -768.46 | 1603.44 | 0.63 |
| CDI production, 24 months | 88 | -1015.73 | 1721.48 | 0.56 |

**Table S15. *Multivariate and univariate linear models for theta cortical tracking at 7 months.*** *Investigating whether theta cortical tracking at 7 months predicted either infant-led or parent-estimated language outcomes. The table details the multivariate linear models (reporting the global effects) and the univariate linear models (ran after the main multivariate model) describing whether theta cortical tracking at 7 months predicted a) infant-led language measures or b) the parent estimated language measures. Bonferroni correction for multiple comparisons in the* ***univariate models*** *led to modified significant alpha levels of, p = <0.01 for infant-led and p = <0.025 for parent-estimated (denoted by* ***bold italic text****).*

### Alpha cortical tracking as a predictor of language acquisition

A multivariate linear model was conducted to see if alpha cortical tracking at 4-, 7- or 11-months predicted either infant-led or parent-estimated language outcomes. Posthoc univariate linear models are also reported in Table S16-S18 to demonstrate the individual contribution of each language measure to global parent-estimated or infant-led language performance.

| **Alpha cortical tracking at 4 months**  **a, Infant-led language measures** |  |  |  |  |
| --- | --- | --- | --- | --- |
| **Multivariate linear model** | *Df _num_* | *Df _denom_* | *F* | *p* |
| Infant-led language measures | 5 | 75 | 1.41 | 0.23 |
| **Univariate linear models (post hoc)** | *Df* | *β _estimate_* | *SE* | *p* |
| NWR Consonants, 24 months | 93 | -6.90 | 7.18 | 0.34 |
| NWR Syllables, 24 months | 93 | -9.27 | 7.06 | 0.19 |
| NWR Stress, 24 months | 93 | -9.62 | 6.65 | 0.15 |
| CCT, 18 months | 92 | -6.28 | 5.13 | 0.22 |
| Pointing, 12 months | 103 | -18.28 | 11.14 | 0.10 |
| **b, Parent-estimated language measures** |  |  |  |  |
| **Multivariate linear model** | *Df _num_* | *Df _denom_* | *F* | *p* |
| Parent-estimated language measures | 2 | 84 | 1.28 | 0.28 |
| **Univariate linear models (post hoc)** | *Df* | *β _estimate_* | *SE* | *p* |
| CDI comprehension, 24 months | 87 | -1621.97 | 3579.18 | 0.65 |
| CDI production, 24 months | 87 | -4845.62 | 3844.16 | 0.21 |

**Table S16. *Multivariate and univariate linear models for alpha cortical tracking at 4 months.*** *Investigating whether alpha cortical tracking at 4 months predicted either infant-led or parent-estimated language outcomes. The table details the multivariate linear models (reporting the global effects) and the univariate linear models (ran after the main multivariate model) describing whether alpha cortical tracking at 4 months predicted a) infant-led language measures or b) the parent estimated language measures. Bonferroni correction for multiple comparisons in the* ***univariate models*** *led to modified significant alpha levels of, p = <0.01 for infant-led and p = <0.025 for parent-estimated (denoted by* ***bold italic text****).*

| **Alpha cortical tracking at 7 months**  **a, Infant-led language measures** |  |  |  |  |
| --- | --- | --- | --- | --- |
| **Multivariate linear model** | *Df _num_* | *Df _denom_* | *F* | *p* |
| Infant-led language measures | 5 | 75 | 0.64 | 0.67 |
| **Univariate linear models (post hoc)** | *Df* | *β _estimate_* | *SE* | *p* |
| NWR Consonants, 24 months | 95 | -7.20 | 6.10 | 0.24 |
| NWR Syllables, 24 months | 95 | -10.66 | 5.95 | 0.08 |
| NWR Stress, 24 months | 95 | -8.76 | 5.62 | 0.12 |
| CCT, 18 months | 90 | 1.69 | 4.72 | 0.72 |
| Pointing, 12 months | 102 | 0.64 | 9.98 | 0.95 |
| **b, Parent-estimated language measures** |  |  |  |  |
| **Multivariate linear model** | *Df _num_* | *Df _denom_* | *F* | *p* |
| Parent-estimated language measures | 2 | 84 | 0.21 | 0.81 |
| **Univariate linear models (post hoc)** | *Df* | *β _estimate_* | *SE* | *p* |
| CDI comprehension, 24 months | 88 | 1886.22 | 3050.86 | 0.54 |
| CDI production, 24 months | 88 | 1260.29 | 3256.13 | 0.70 |

**Table S17. *Multivariate and univariate linear models for alpha cortical tracking at 7 months.*** *Investigating whether alpha cortical tracking at 7 months predicted either infant-led or parent-estimated language outcomes. The table details the multivariate linear models (reporting the global effects) and the univariate linear models (ran after the main multivariate model) describing whether alpha cortical tracking at 7 months predicted a) infant-led language measures or b) the parent estimated language measures. Bonferroni correction for multiple comparisons in the* ***univariate models*** *led to modified significant alpha levels of, p = <0.01 for infant-led and p = <0.025 for parent-estimated (denoted by* ***bold italic text****).*

| **Alpha cortical tracking at 11months**  **a, Infant-led language measures** |  |  |  |  |
| --- | --- | --- | --- | --- |
| **Multivariate linear model** | *Df _num_* | *Df _denom_* | *F* | *p* |
| Infant-led language measures | 5 | 77 | 0.89 | 0.49 |
| **Univariate linear models (post hoc)** | *Df* | *β _estimate_* | *SE* | *p* |
| NWR Consonants, 24 months | 96 | -3.59 | 6.60 | 0.59 |
| NWR Syllables, 24 months | 96 | -8.60 | 6.55 | 0.19 |
| NWR Stress, 24 months | 96 | -0.37 | 6.03 | 0.95 |
| CCT, 18 months | 93 | 1.79 | 5.24 | 0.73 |
| Pointing, 12 months | 105 | 2.75 | 10.55 | 0.79 |
| **b, Parent-estimated language measures** |  |  |  |  |
| **Multivariate linear model** | *Df _num_* | *Df _denom_* | *F* | *p* |
| Parent-estimated language measures | 2 | 87 | 0.44 | 0.65 |
| **Univariate linear models (post hoc)** | *Df* | *β _estimate_* | *SE* | *p* |
| CDI comprehension, 24 months | 90 | -3122.35 | 3336.18 | 0.35 |
| CDI production, 24 months | 90 | -2955.11 | 3613.68 | 0.42 |

**Table S18. *Multivariate and univariate linear models for alpha cortical tracking at 11 months.*** *Investigating whether alpha cortical tracking at 11 months predicted either infant-led or parent-estimated language outcomes. The table details the multivariate linear models (reporting the global effects) and the univariate linear models (ran after the main multivariate model) describing whether alpha cortical tracking at 11 months predicted a) infant-led language measures or b) the parent estimated language measures. Bonferroni correction for multiple comparisons in the* ***univariate models*** *led to modified significant alpha levels of, p = <0.01 for infant-led and p = <0.025 for parent-estimated (denoted by* ***bold italic text****).*

### PAC as a predictor of language acquisition

A multivariate linear model was conducted to see if PAC at 7- or 11-months predicted either infant-led or parent-estimated language outcomes. Posthoc univariate linear models are also reported in Table S19 & S20 to demonstrate the individual contribution of each language measure to global parent-estimated or infant-led language performance.

| **PAC at 7 months**  **a, Infant-led language** |  | | | |  | | |  | |  | |
| --- | --- | --- | --- | --- | --- | --- | --- | --- | --- | --- | --- |
|  | *Delta-beta* | | *Delta-gamma* | | | *Theta-beta* | | | *Theta-gamma* | | |
| **Multivariate linear model** | *F* | *p* | *F* | *p* | | *F* | *p* | | *F* | | *p* |
| Infant-led language measures | 0.73 | 0.61 | 2.19 | **0.06** | | 0.31 | 0.90 | | 1.17 | | 0.33 |
|  | *Delta-beta* | | *Delta-gamma* | | | *Theta-beta* | | | *Theta-gamma* | | |
| **Univariate linear models** | *β _est._* | *p* | *β _est._* | *p* | | *β _est._* | *p* | | *β _est._* | | *p* |
| NWR Consonants, 24 months | 0.03 | 0.64 | -0.02 | 0.76 | | -0.07 | 0.27 | | -0.05 | | 0.37 |
| NWR Syllables, 24 months | 0.08 | 0.19 | -0.02 | 0.81 | | -0.04 | 0.55 | | -0.05 | | 0.28 |
| NWR Stress, 24 months | -0.05 | 0.46 | -0.13 | 0.04 | | -0.10 | 0.07 | | -0.04 | | 0.38 |
| CCT, 18 months | -0.00 | 0.94 | -0.04 | 0.58 | | -0.03 | 0.57 | | -0.02 | | 0.67 |
| Pointing, 12 months | -0.02 | 0.88 | 0.05 | 0.67 | | 0.06 | 0.55 | | 0.04 | | 0.60 |
| **b, Parent-estimated language** |  | | | |  | | |  | |  | |
|  | *Delta-beta* | | *Delta-gamma* | | | *Theta-beta* | | | *Theta-gamma* | | |
| **Multivariate linear model** | *F* | *p* | *F* | *p* | | *F* | *p* | | *F* | | *p* |
| Parent-estimated lang measures | 2.16 | 0.12 | 0.02 | 0.98 | | 2.37 | **.099** | | 1.48 | | 0.23 |
|  | *Delta-beta* | | *Delta-gamma* | | | *Theta-beta* | | | *Theta-gamma* | | |
| **Univariate linear models** | *β _est._* | *p* | *β _est._* | *p* | | *β _est._* | *p* | | *β _est._* | | *p* |
| CDI comprehension, 24 mo | -5.57 | 0.86 | 0.41 | 0.99 | | -55.73 | 0.08 | | -35.59 | | 0.18 |
| CDI production, 24 mo | 36.00 | 0.30 | 4.27 | 0.91 | | -25.81 | 0.45 | | -14.24 | | 0.62 |

**Table S19. Multivariate and univariate linear models for PAC at 7 months.** Investigating whether delta-beta, delta-gamma, theta-beta or theta-gamma PAC at 7 months predicted either infant-led or parent-estimated language outcomes. The table details the multivariate linear models (reporting the global effects) and the univariate linear models (ran after the main multivariate model) describing whether PAC at 7 months predicted a) infant-led language measures or b) the parent-estimated language measures. Bonferroni correction for multiple comparisons in the **univariate models** led to modified significant alpha levels of, p = <0.01 for infant-led and p = <0.025 for parent-estimated (denoted by bold italic text). Modified alpha level for trends are p = <0.02 for infant-led and p = <0.05 for parent-estimated (denoted by bold text).

| **PAC at 11 months**  **a, Infant-led language** |  | | | |  | | |  | |  | |
| --- | --- | --- | --- | --- | --- | --- | --- | --- | --- | --- | --- |
|  | *Delta-beta* | | *Delta-gamma* | | | *Theta-beta* | | | *Theta-gamma* | | |
| **Multivariate linear model** | *F* | *p* | *F* | *p* | | *F* | *p* | | *F* | | *p* |
| Infant-led language measures | 1.10 | 0.35 | 0.47 | 0.80 | | 1.20 | 0.32 | | 0.17 | | 0.97 |
|  | *Delta-beta* | | *Delta-gamma* | | | *Theta-beta* | | | *Theta-gamma* | | |
| **Univariate linear models** | *β _est._* | *p* | *β _est._* | *p* | | *β _est._* | *p* | | *β _est._* | | *p* |
| NWR Consonants, 24 months | 0.19 | 0.02 | 0.05 | 0.58 | | 0.07 | 0.23 | | -0.02 | | 0.79 |
| NWR Syllables, 24 months | 0.19 | **.0197** | 0.08 | 0.32 | | 0.12 | 0.04 | | 0.01 | | 0.83 |
| NWR Stress, 24 months | 0.09 | 0.25 | 0.02 | 0.78 | | 0.04 | 0.48 | | 0.01 | | 0.87 |
| CCT, 18 months | 0.10 | 0.12 | -0.04 | 0.53 | | -0.01 | 0.87 | | -0.06 | | 0.25 |
| Pointing, 12 months | -0.02 | 0.88 | 0.05 | 0.73 | | 0.05 | 0.63 | | -0.01 | | 0.93 |
| **b, Parent-estimated language** |  | | | |  | | |  | |  | |
|  | *Delta-beta* | | *Delta-gamma* | | | *Theta-beta* | | | *Theta-gamma* | | |
| **Multivariate linear model** | *F* | *p* | *F* | *p* | | *F* | *p* | | *F* | | *p* |
| Parent-estimated lang measures | 1.68 | 0.19 | 0.13 | 0.88 | | 0.99 | 0.37 | | 2.23 | | 0.11 |
|  | *Delta-beta* | | *Delta-gamma* | | | *Theta-beta* | | | *Theta-gamma* | | |
| **Univariate linear models** | *β _est._* | *p* | *β _est._* | *p* | | *β _est._* | *p* | | *β _est._* | | *p* |
| CDI comprehension, 24 mo | 56.88 | 0.17 | -7.05 | 0.87 | | 0.52 | 0.99 | | -66.61 | | **0.04** |
| CDI production, 24 mo | 80.33 | 0.07 | 6.00 | 0.89 | | 26.78 | 0.42 | | -58.58 | | 0.09 |

**Table S20. Multivariate and univariate linear models for PAC at 11 months**. Investigating whether delta-beta, delta-gamma, theta-beta or theta-gamma PAC at 11 months predicted either infant-led or parent-estimated language outcomes. The table details the multivariate linear models (reporting the global effects) and the univariate linear models (ran after the main multivariate model) describing whether PAC at 11 months predicted a) infant-led language measures or b) the parent-estimated language measures. Bonferroni correction for multiple comparisons in the **univariate models** led to modified significant alpha levels of, p = <0.01 for infant-led and p = <0.025 for parent-estimated (denoted by bold italic text). Modified alpha level for trends are p = <0.02 for infant-led and p = <0.05 for parent-estimated (denoted by bold text).
